# A transcriptomics-based drug repositioning approach to identify drugs with similar activities for the treatment of muscle pathologies in spinal muscular atrophy (SMA) models

**DOI:** 10.1101/2023.06.14.544899

**Authors:** Joseph M Hoolachan, Eve McCallion, Emma R Sutton, Özge Çetin, Paloma Pacheco-Torres, Maria Dimitriadi, Magnus Okoh, Lisa M Walter, Peter Claus, Matthew JA Wood, Daniel P Tonge, Melissa Bowerman

## Abstract

Spinal muscular atrophy (SMA) is a genetic neuromuscular disorder caused by the reduction of survival of motor neuron (SMN) protein levels. Although three SMN-augmentation therapies are clinically approved that significantly slow down disease progression, they are unfortunately not cures. Thus, complementary SMN-independent therapies that can target key SMA pathologies and that can support the clinically approved SMN-dependent drugs are the forefront of therapeutic development. We have previously demonstrated that prednisolone, a synthetic glucocorticoid (GC) improved muscle health and survival in severe *Smn^-/-^;SMN2* and intermediate *Smn^2B/-^* SMA mice. However, long-term administration of prednisolone can promote myopathy. We thus wanted to identify genes and pathways targeted by prednisolone in skeletal muscle to discover clinically approved drugs that are predicted to emulate prednisolone’s activities. Using an RNA-sequencing, bioinformatics and drug repositioning pipeline on skeletal muscle from symptomatic prednisolone-treated and untreated *Smn^-/-^;SMN2* SMA and *Smn^+/-^;SMN2* healthy mice, we identified molecular targets linked to prednisolone’s ameliorative effects and a list of 580 drug candidates with similar predicted activities. Two of these candidates, metformin and oxandrolone, were further investigated in SMA cellular and animal models, which highlighted that these compounds do not have the same ameliorative effects on SMA phenotypes as prednisolone; however, a number of other important drug targets remain. Overall, our work further supports the usefulness of prednisolone’s potential as a second-generation therapy for SMA, identifies a list of potential SMA drug treatments and highlights improvements for future transcriptomic-based drug repositioning studies in SMA.

## INTRODUCTION

Spinal muscular atrophy (SMA) is a heterogenous autosomal recessive neuromuscular disorder (NMD) characterized by motor neuron degeneration alongside progressive muscle atrophy and weakness ^1^. Being the leading monogenic cause of infant mortality ^2^, around 96% of SMA cases are mapped to homozygous loss-of-function and deletion mutations in the *survival of motor neuron 1* (*SMN1*) gene ^3, 4^, which ubiquitously expresses SMN, a protein that current and ongoing research has linked to diverse housekeeping and tissue-specific cellular functions ^5–7^. Although complete SMN loss is embryonic lethal in most organisms^8^, humans can overcome the complete loss of the *SMN1* gene due to incomplete rescue by the homologous *SMN2* gene ^9, 10^. In essence, the presence of a single nucleotide mutation in *SMN2* promotes exon 7 alternative splicing that limits full length SMN (FL-SMN) expression in this gene to 10% ^11^. Consequently, the limited FL-SMN expression makes *SMN2* gene copy number an important disease modifier, impacting SMA type and severity^12^.

In recent years, novel SMN restorative SMA treatments have emerged that either increase FL- *SMN2* expression by an anti-sense oligonucleotide (ASO) (Nusinersen marketed as (Spinraza®)) ^13, 14^ or a small molecule (Evrysdi®) ^15, 16^ or promote exogenous FL-*SMN1* expression by an adeno-associated virus 9 (AAV-9) delivery system (Zolgensma®) ^17, 18^. Despite the significant increased life expectancy and improved quality of life associated with these therapies ^14, 16, 18, 19^, they are not cures and their efficacy is dependent upon early intervention ^20^.

Thus, additional SMN-independent therapies that target affected tissues such as muscle are needed to further enhance and support the benefits of SMN-dependent treatments ^21^. Indeed, pre-clinical studies and primary patient data have reported innate muscular defects in SMA, which include myogenesis ^22^, regeneration ^23^, contraction ^24, 25^, regulation ^25^, growth ^26^, and metabolism ^27^, highlighting skeletal muscle as a primary therapeutic target. Although two novel skeletal muscle-specific SMA therapies, Apitegromab^TM^ ^28^ (ClinicalTrials.gov ID: NCT03921528) and Reldesemtiv^TM^ ^29^ (ClinicalTrials.gov ID: NCT02644668), are showing progress in clinical trials, the high expenses involved in novel drug research and development (R&D) ^30^ alongside the costs of the three clinically approved SMN-dependent therapies may lead to elevated prices for combinatorial treatments ^31^, thus further widening the accessibility gap for SMA patients.

A useful alternative for ensuring accessibility of SMN-independent treatments for all SMA patients would be the identification of cost-effective generic drugs via drug repositioning, a strategy aimed at finding new therapeutic activities for existing pharmacological compounds ^32^. One such example is prednisolone, a synthetic glucocorticoid (GC) administered to relieve muscle inflammation in Duchenne muscular dystrophy (DMD) patients ^33, 34^. Interestingly, evidence also emerged that short-term prednisolone treatment additionally conferred ergogenic muscle benefits in DMD patients ^35, 36^, which was also validated in the *mdx* mouse model of DMD ^37, 38^.

Although GCs are not used for the treatment of SMA patients, prednisolone is administered for a short period (∼30 days, 1 mg/kg) to alleviate the immunological adverse effects of Zolgensma®^18^. However, given prednisolone’s potential muscle benefits, we have previously investigated and demonstrated that prednisolone treatment (5 mg/kg, every other day) in severe *Smn^-/-^;SMN2* and intermediate *Smn^2B/-^* SMA mice improved survival, weight and neuromuscular phenotype ^27^. Although the study was aimed at investigating prednisolone’s activity on the GC-Krüppel-Like-Factor-15 (KLF15) pathway in SMA skeletal muscle, synergistic muscle improvement was also observed in prednisolone-treated *Smn^-/-^;SMN2* SMA mice overexpressing Klf15 specifically in skeletal muscle ^27^, suggesting that prednisolone may act on SMA skeletal muscle via numerous effectors and pathways.

Despite the benefits observed, our study did not evaluate prednisolone’s long-term effects ^18, 27^. Given that chronic GC usage increases myopathy ^35, 39^, it is unclear whether long-term prednisolone treatments would be similarly detrimental in SMA muscle. Furthermore, the rapid onset and progression of disease in SMA mouse models (1-3 weeks on average) ^9, 40, 41^ does not allow sufficient comparison of intermittent vs chronic studies.

Thus, in this study, we used a transcriptomics and drug repositioning pipeline based on our previously published experimental paradigm ^42^ to uncover the genes and pathways restored by prednisolone in skeletal muscle of SMA mice and identify existing non-GC drugs predicted to have similar activities. Our study uncovered that prednisolone restored pathways linked to growth, metabolism, and regulation in SMA skeletal muscle and identified 20 leading commercially available non-GC drugs predicted to emulate its action. Based on oral bioavailability and evidence of safety treatment in children, we selected and validated metformin and oxandrolone in SMA cellular and animal models. Although, both metformin and oxandrolone improved neuromuscular activity in the *Caenorhabditis elegans (C. elegans)* model for severe SMA, we found that higher metformin doses reduced survival in the intermediate *Smn^2B/-^* SMA mouse model. On the other hand, oxandrolone treatment partially improved survival in *Smn^2B/-^* SMA mice, albeit not to the same extent as prednisolone ^27^.

Nevertheless, our study computationally uncovered new mechanisms behind prednisolone’s beneficial activity in SMA muscle, identified numerous potential SMA muscle-specific therapeutic candidates and highlighted the importance of transcriptomic-based drug repositioning for SMN-independent drug discovery.

## METHODS

### Animal Procedures

Experiments with the severe (Taiwanese) *Smn^-/-^;SMN2 (*FVB/N background, FVB.Cg- *Smn1tm1HungTg(SMN2)2Hung/J*) SMA mice ^40^ and *Smn^+/-^;SMN2* healthy control littermates were carried out in the University of Oxford Biomedical Sciences Unit (BSU), in accordance with the UK Home Office authorisation (Animals Scientific Procedures Act (1986), UK Home Office Project Licence PDFEDC6F0).

The *Smn^2B/-^* SMA mouse model ^43^ was generated by breeding *Smn^2B/2B^* mice (generously provided by Dr Rashmi Kothary (University of Ottawa), Dr Lyndsay Murray (University of Edinburgh) and Professor Matthew Wood (University of Oxford) before being sent to Charles River for rederivation) with *Smn^+/-^* mice (B6.Cg-*Smn1/J*, stock #007963, Jackson Labs). All live procedures on wild type (WT) (C57BL/6 background), *Smn^2B/-^* SMA and *Smn^2B/+^* healthy littermates were performed in the Keele University BSU, in accordance with the UK Home Office authorisation (Animals Scientific Procedures Act (1986), UK Home Office Project Licence P99AB3B95).

For all behavioural experiments, body weights and righting reflex ^44^ (up to 30 seconds) were assessed daily from birth until animals reached their humane endpoint, defined in our UK Home Office Project Licence (P99AB3B95) as the time at which the animal displays hindlimb paralysis, immobility, inability to right (greater than 30 seconds), 4 consecutive days of weight loss and/or up to 20% body weight loss.

As previously described ^27^, prednisolone (5 mg tablet, Almus) (5 mg/kg dissolved in water) was administered every second day by gavage from post-natal day (P)0 to P7 in *Smn^-/-^;SMN2* SMA and *Smn^+/-^;SMN2* healthy mice and from P0 to P20 in *Smn^2B/-^* SMA and *Smn^2B/+^* healthy mice. Metformin hydrochloride (#PHR1084, Sigma-Aldrich) was dissolved in 0.9% saline physiological solution (tablet dissolved in sterile water) (#07982, Sigma) and administered daily (200 or 400 mg/kg) by gavage from P5 to humane endpoint in *Smn^2B/-^* SMA, *Smn^2B/+^* healthy and WT mice. Oxandrolone (#SML0437, Sigma-Aldrich) was prepared in 0.5% carboxymethylcellulose (CMC) solution (powder dissolved in 0.9% saline solution) (#C5678, SLS) by sonication at 37 kHz for 3 minutes and administered (4 mg/kg) daily by gavage from P8 to P21 in *Smn^2B/-^* SMA, *Smn^2B/+^* healthy and WT mice.

### Blood-glucose measurement

Blood was collected from non-fasted *Smn^2B/-^* SMA and *Smn^2B/+^* healthy mice and glucose levels were immediately measured (mmol/L) via True Metrix® GO Self-Monitoring Blood Glucose Meter (Trividia Health^TM^) and True Metrix® Test Strips (Trividia Health^TM^).

### RNA-Sequencing (RNA-Seq)

Total RNA was extracted from triceps of symptomatic P7 untreated and prednisolone-treated Taiwanese *Smn^-/-^;SMN2* SMA and *Smn^+/-^;SMN2* healthy mice (Table S1). The triceps were immediately placed in RNALater (#AM7030, ThermoFisher) following dissection and stored at - 20°C under further analysis. For mRNA isolation, 500 ng of total RNA was used as input for the NEBNext® Poly(A) mRNA Magnetic Isolation Module’ (#E7490L, New England Biolabs (NEB)) in accordance with the manufacturer’s standard instruction. Library preparation was carried out using the NEBNext® Ultra Directional RNA Library Prep Kit for Illumina (#E7420L, NEB). Barcoded libraries from each experimental sample were combined in equimolar concentrations of 1.5 pM prior to sequencing at 75bp x 1 (single-end) read metric on a NextSeq 550 (Illumina) system.

### Differential gene expression analysis

For the RNA-Seq data from the triceps of P7 untreated and prednisolone-treated Taiwanese *Smn^-/-^;SMN2* SMA and *Smn^+/-^;SMN2* healthy mice (Table S1), differential gene expression (DGE) analysis was performed in Galaxy (usegalaxy.org) ^45^. After initial quality control assessments via FastQC v0.72+galaxy1 ^46^, we trimmed reads based on SLIDINGWINDOW of 4 bp at average quality read of 32 in Trimmomatic v0.36.5 ^47^ and trimmed the first 12 abnormal bases in Trim sequences v1.0.2 ^48^. After quality control confirmation with FastQC v0.72+galaxy1 ^46^ the processed 63 bp single-reads were aligned to an in-built UCSC *Mus musculus* mm10 genome via HISAT2 v2.1.0 ^45, 49^ under a reverse (or antisense) strand setting. Count quantification of aligned single-reads to mapped coding genes was performed by FeatureCounts v1.6.3+galaxy2 ^45, 50^ using an in-built Entrez *Mus musculus* mm10 gene transfer file (GTF) with known gene identifier set at*-exon level*. Mapping and count quantification was visualized through MultiQC v1.6 ^51^. For DGE analysis of our raw transcript counts, we used DESeq2 v2.11.40.2 ^52^ under the design formula of “Condition” (SMA vs Healthy) and “Treatment” (Untreated vs Prednisolone) after removal of 1 outlier (prednisolone-treated *Smn^-/-^;SMN2* sample N0603). We set differentially expressed gene (DEG) significance at log2 fold change (FC) > 0.6 and false discovery rate (FDR) < 0.05. The Entrez Gene IDs were translated to official *Mus musculus* gene symbols by AnnotatemyID v3.7.0^45^. The normalized count files generated by DESeq2 of prednisolone-treated Taiwanese *Smn^-/-^ ;SMN2* SMA mice vs untreated *Smn^-/-^;SMN2* SMA and *Smn^+/-^;SMN2* healthy mice were generated into heatmaps by Heatmap2 v2.2.1+galaxy1 ^45^ under -*data log2 transformed* and scaled by *-row scale genes*.

### Pathway analysis and drug repositioning

Pathway analysis of the prednisolone-treated vs untreated Taiwanese *Smn^-/-^;SMN2* SMA mice was performed with iPathwayGuide ^53^ (Advaita) with default criteria of log2FC > 0.6 and FDR < 0.05 for DEGs. The impact analysis performed by iPathwayGuide incorporated our DEGs expression (log2FC) and its topological position in the KEGG pathway database ^54^ to calculate significantly impacted pathways *p* < 0.05 (KEGG v1910 Release 90.0+/05-29, May 19, GODb v1910 2019- Apr 26). Furthermore, overrepresentation analysis ^55^ (ORA) with elimination pruning ^56^ was performed for gene ontology (GO) pathways ^57^ and predicted upstream regulators (STRING v11.0, Jan 19^th^, 2019). Drug candidate identification was performed through an in-built KEGG drugs database ^54^ aligned to KEGG pathways in iPathwayGuide and by drug-gene interactions for upstream regulators in the Drug Gene Interaction Database ^58^ (DGIdb) v3.0.

### C2C12 cell culture

The immortalised murine C2C12 myoblast-like cell line ^59^ (#CRL-1772, ATCC, USA) was maintained in growth media comprised of high glucose (4.5 g/L) and L-glutamine (0.6 g/L) Dulbecco’s Modified Eagle’s Media (DMEM) (#BE12-741F, Lonza), 10% foetal bovine serum (FBS) (#10438026, Gibco), and 1% penicillin-streptomycin (10,000 U/ml) (#15140122, Gibco). C2C12 myoblasts were differentiated into myotubes in a differentiation media comprised of high glucose (4.5 g/L) and L-glutamine (0.6 g/L) DMEM (#BE12-741F, Lonza), 2% horse serum (HS) (#26050070, Gibco), 1% penicillin-streptomycin (10,000 U/ml) (#15140122, Gibco), and 0.1% insulin (1 μg/ml) (#I6634, Sigma) for 2–8 days with media replacement every 48 hours. All cultured cells were incubated in humid 37°C and 5% CO2 conditions (Heracell 150i CO2 incubator, ThermoScientific).

### *In vitro* drug treatment

Proliferating C2C12 myoblasts were seeded in 6-well plates (x4 wells per group). *In vitro* drug treatments began at 50-60% confluence for C2C12 myoblasts and D7 stage for C2C12 myotubes. For metformin groups, they were treated with metformin hydrochloride (#PHR1084, Sigma- Aldrich) dissolved in phosphate buffered saline (PBS), pH 7.4 (#10010023, ThermoFisher) at concentrations of 0.3, 0.6, 1 and 2 mM for 24 hours against a PBS control (0.1% v/v). For oxandrolone groups, they were treated with oxandrolone (#SML0437, Sigma-Aldrich) dissolved in ethanol absolute > 99.8% (#20821.296, VWR) at concentrations of 1, 10 and 100 μM for 24 hours against an ethanol absolute > 99.8% vehicle control (0.1% v/v).

### Lactate dehydrogenase (LDH) assay

Drug cytotoxicity was measured by the lactate dehydrogenase (LDH)-Glo^TM^ Cytotoxicity Assay Kit (#J2380, Promega) following manufacturer’s instructions. Luminescence was measured at 400 nm using the GloMax® Explorer Multimode Microplate Reader (#GM3500, Promega).

### Bromodeoxyuridine (BrDU) cell proliferation assay

Cell proliferation was measured by the Bromodeoxyuridine (BrDU) Cell Proliferation Assay Kit (#QIA58, Sigma-Aldrich) following manufacturer’s instructions. Absorbance was measured at 450 – 540 nm using the GloMax® Explorer Multimode Microplate Reader (#GM3500, Promega).

### Small interfering RNA-mediated *Smn* knockdown in C2C12 cells

A 10 μM *Smn* small interfering RNA (siRNA) (Duplex name: mm.RiSmn1.13.1) (Integrated DNA Technologies (IDT)) was used to knock down *Smn* levels against a 10 μM scrambled siRNA (scrambled DsiRNA, #51-01-19-08) (IDT) negative control. The *Smn* and scrambled siRNAs were aliquoted separately into an siRNA-lipofectamine complex containing Lipofectamine® RNAiMAX Reagent (#13778075, Invitrogen) and Opti-MEM (#31985062, Gibco) following manufacturer’s instructions. C2C12 myoblasts were transfected for 48 hours with *Smn* depletion, whilst C2C12 myotubes were freshly transfected at differentiation (D) stages D0, D3 and D6 for 48 hours with *Smn* depletion confirmed via quantitative polymerase chain reaction (qPCR) (Table S2).

### Serum starvation-induced canonical muscle atrophy in differentiated C2C12 myotubes

Differentiated C2C12 myotubes were incubated in serum-free glucose (4.5 g/L) and L-glutamine (0.6 g/L) DMEM (#BE12-741F, Lonza) with 1% Penicillin-Streptomycin (10,000 U/ml) (#15140122, Gibco) for 24 hours. Atrophy was confirmed by upregulation of atrogenes *Atrogin-1* and *MuRF1* via qPCR (Table S2) and morphology via microscopy (Motic AE31E).

### RNA extraction and quantitative polymerase chain reaction (qPCR)

RNA extraction for C2C12 cells was performed with the ISOLATE II RNA Mini Kit (#BIO- 52073, Meridian BIOSCIENCE) as per manufacturer’s instructions. Skeletal muscle (triceps and *Tibialis anterior* (TA)) and spinal cord tissue samples underwent homogenization with 7 mm stainless steel beads (#69990, Qiagen) in a Tissue Lyser LT (#85600, Qiagen) set at 60 oscillations/second for 2 minutes followed by microcentrifugation at >10,000 RCF (MSE Sanyo Hawk 15/05) for 1 minute. RNA extractions from harvested skeletal muscle was performed with the RNeasy Fibrous Tissue Kit (#74704, Qiagen) and all other harvested tissues with ISOLATE II RNA Mini Kit (#BIO-52073, Meridian BIOSCIENCE) as per manufacturer’s instructions. RNA concentrations (ng/μl) were measured with the NanoDrop 1000 spectrophotometer (ThermoScientific) before reverse transcription was performed using the qPCRBIO cDNA Synthesis Kit (#PB30.11-10, PCR Biosystems) as per manufacturer’s instructions.

The cDNA was then diluted by 1:5 in nuclease-free water (#10526945, ThermoFisher). qPCR was performed using 2x PCRBIO Sygreen Blue Mix Hi-ROX (#PB20.16-20, PCR Biosystems), nuclease-free water (#10526945, ThermoFisher) and 10 μM forward and reverse primers obtained from IDT (Table S2). The qPCR reactions were performed in the StepOnePlus^TM^ Real-Time PCR System (ThermoScientific) with the following program: initial denaturation (95°C for 2 minutes), 40 cycles of 95 °C for 5 seconds and 60 °C for 30 seconds and ending with melt curve stage (95 °C for 15 seconds, 60 °C for 1 minutes and 95 °C for 15 seconds). Relative gene expression was quantified using the Pfaffl method ^60^ and referenced against the validated *RNA polymerase II polypeptide J* (*PolJ)* housekeeping gene ^27, 61^ (Table S2). Primer efficiency was calculated using LinRegPCR V11.0 ^62^.

### Caenorhabditis elegans drug treatments

The *Caenorhabditis elegans* (*C. elegans*) SMA strains used in this study was LM99 *smn- 1(ok355)I/hT2(I;III)*, which was segregated into homozygotes *smn-1(ok355)*, lethal homozygotes *hT2/hT2* and control heterozygotes *smn-1(ok355)I/hT2* ^63^. These animals were maintained at 20°C on Nematode Growth Medium (NGM) plates seeded with *Escherichia coli* OP50 bacteria ^64^. *C. elegans* were treated by mixing metformin hydrochloride (#PHR1084, Sigma-Aldrich) dissolved in water at concentrations of 0, 1, 10 and 50 mM and oxandrolone (#SML0437, Sigma-Aldrich) dissolved in DMSO at concentrations of 0, 1, 10 and 50 μM with NGM agar solution.

### Caenorhabditis elegans neuromuscular assays

Neuromuscular assays were performed on day 3 animals raised on plates containing the pertinent solvent or drug. The pharyngeal pumping assay was performed as previously described ^65^. Briefly, animals were filmed with a 150x objective using a AxioCam ICc5 camera at 175 frames/10sec on a Discovery.V8 SteREO microscope. Pumps were manually counted using the Zen Pro software v2.3. A pumping event was defined as a grinder movement in any axis. For locomotion assays animals were filmed with a 63x objective using a AxioCam ICc5 camara at 15 frames/sec on a Discovery.V8 SteREO microscope. Reversals, and paralysis time for 5 minutes (±SEM) were quantified using WormLab 1.1 software (MBF Bioscience). Final data represents three independent trials (n ≥ 25 animals in total per genotype).

### Statistical Analyses

Statistical analyses were carried out using the most up to date GraphPad PRISM software. Prior to any analyses, outliers were identified via Grubb’s test (GraphPad) and subsequently removed. Appropriate statistical tests include unpaired t-test, one-way analysis of variance (ANOVA), and two-way ANOVA. Each post-hoc analyses used is noted in the respective figure legend. Kaplan-Meier survival curves were analysed with a log-rank test. Statistical significance was considered at *p* < 0.05, described in graphs as \**p* < 0.05, \*\**p* < 0.01, \*\*\**p* < 0.001 and \*\*\*\**p* < 0.0001.

## RESULTS

### Prednisolone restores the expression of a large subset of genes involved in canonical skeletal muscle pathways in SMA mice

As described in an earlier study, we have previously demonstrated that treating SMA mice with prednisolone significantly improved several disease phenotypes, including survival, weight, and muscle health ^27^. To have a more in depth understanding of the impact of prednisolone on SMA skeletal muscle at a molecular level, we performed bulk RNA-Seq on skeletal muscle of untreated and prednisolone-treated *Smn^-/-^;SMN2* SMA and *Smn^+/-^;SMN2* healthy mice. Specifically, we administered prednisolone (5 mg/kg, gavage, every 2 days) starting from P0 until P7 to *Smn^-/-^;SMN2* SMA and *Smn^+/-^;SMN2* healthy mice ^27^. Triceps were harvested from P7 prednisolone-treated and untreated mice for RNA-Seq via Illumina NextSeq550 and a HISAT2-FeatureCounts- DESeq2 pipeline against a Mus Musculus mm10 genome for parameters of “condition” and “treatment” (Figures S1).

Initially, our principal component analysis (PCA) revealed distinct clusters of untreated *Smn^-/-^;SMN2* SMA and untreated *Smn^+/-^;SMN2* healthy littermates, with prednisolone-treated *Smn^-/-^;SMN2* SMA mice falling between the aforementioned groups (Figure 1.a). Importantly, we found that prednisolone treatment restored the expression of 1361 genes in *Smn^-/-^;SMN2* SMA mice to levels similarly observed in untreated *Smn^+/-^;SMN2* healthy mice (Figure 1.b; Tables S3-5).

**Figure 1.**
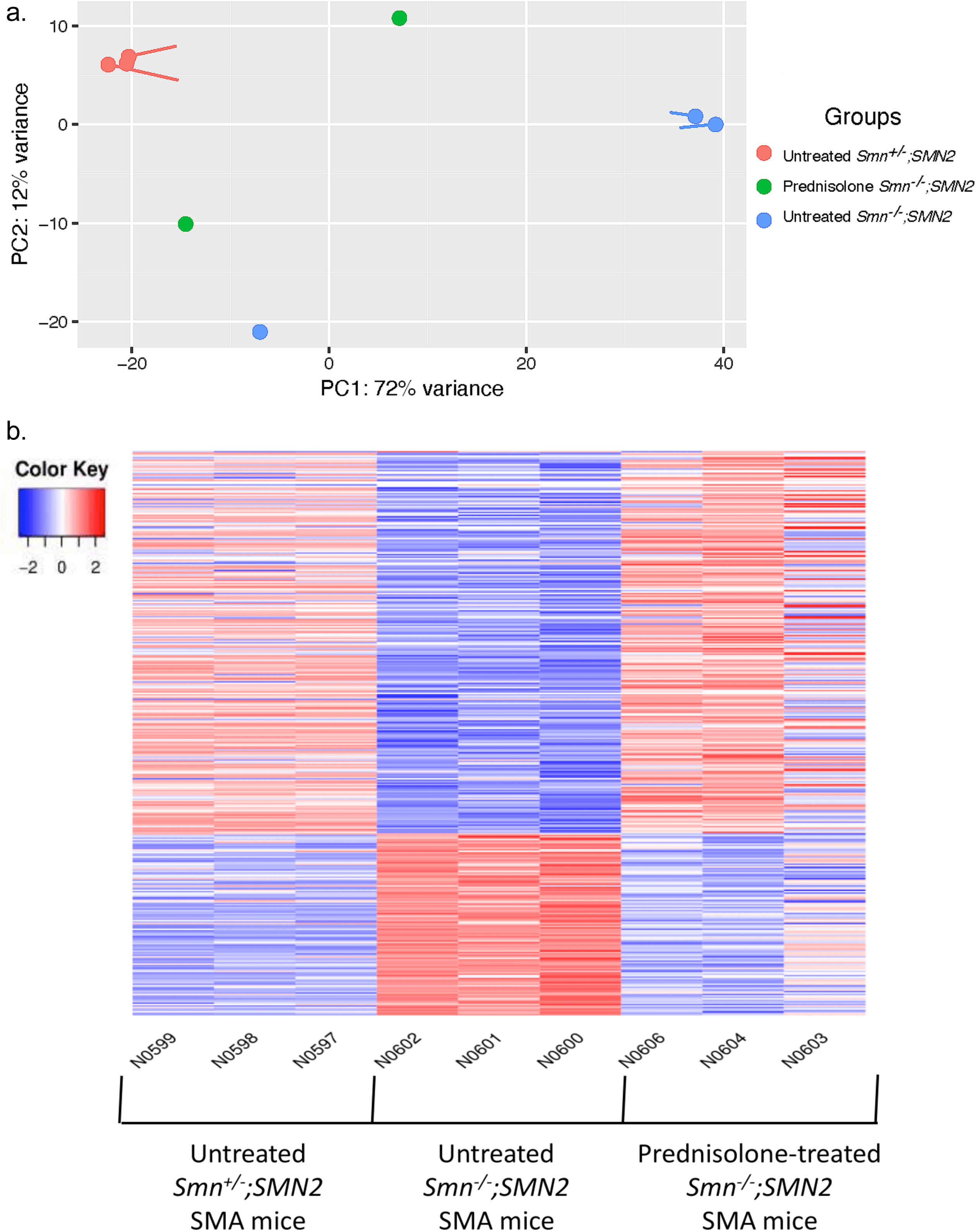
Prednisolone treatment normalizes a subset of genes in severe *Smn^-/-^;SMN2* SMA mice to healthy levels observed in untreated *Smn^+/-^;SMN2* mice. *Smn^-/-^;SMN2* SMA and *Smn^+/-^;SMN2* healthy mice received prednisolone treatment (5 mg/kg gavage every 2 days) from P0. The triceps was harvested from P7 untreated and prednisolone- treated *Smn^-/-^;SMN2* SMA and *Smn^+/-^;SMN2* healthy mice for RNA isolation and library preparation for RNA-Sequencing. Differential gene expression analysis was performed by DESeq2 v2.11.40.2 with study design set to “condition and “treatment”. (**a)** Principal component analysis based on transcriptomic profiles between P7 untreated *Smn^+/-^;SMN2* (red, n=3), prednisolone-treated *Smn^-/-^;SMN2* (green, n=2) and untreated *Smn^-/-^;SMN2* (blue, n=3) mice. **(b)** Heatmap of the transcriptomic expression profiles (Log2FC >0.6; FDR <0.05) between P7 untreated *Smn^+/-^;SMN2* (n=3, left), untreated *Smn^-/-^;SMN2* (n=3, centre) and prednisolone-treated *Smn^-/-^;SMN2* (n=3, right) mice. Upregulated genes highlighted in red and downregulated genes highlighted in blue.

Next, we determined the biological pathways associated with DEGs in prednisolone-treated *Smn^-/-^;SMN2* SMA mice compared to untreated *Smn^-/-^;SMN2* SMA mice. Using iPathwayGuide, we identified that 3056 significant DEGs (Log2FC > 0.6, FDR < 0.05) (Table S4) were targeted by prednisolone in the skeletal muscle of *Smn^-/-^;SMN2* SMA mice when compared to untreated *Smn^-/-^;SMN2* SMA mice and associated with 28 significant KEGG pathways (*p* <0.05) (Table 1).

**Table 1.** KEGG pathways targeted in the skeletal muscle of symptomatic prednisolone- treated *Smn^-/-^;SMN2* SMA mice compared with untreated *Smn^-/-^;SMN2* SMA mice.

| KEGG ID | Pathway name | #Genes (DE/All) | p value |
| --- | --- | --- | --- |
| 04066 | HIF-1 signaling pathway | 30/91 | 0.001 |
| 05214 | Glioma | 25/71 | 0.002 |
| 04213 | Longevity regulating pathway - multiple species | 24/57 | 0.003 |
| 04068 | FoxO signaling pathway | 41/116 | 0.003 |
| 04713 | Circadian entrainment | 23/73 | 0.004 |
| 04211 | Longevity regulating pathway | 30/83 | 0.004 |
| 04115 | p53 signaling pathway | 25/67 | 0.006 |
| 00040 | Pentose and glucuronate interconversions* | 8/15 | 0.007 |
| 05010 | Alzheimer disease | 19/158 | 0.008 |
| 04152 | AMPK signaling pathway | 37/110 | 0.008 |
| 04080 | Neuroactive ligand-receptor interaction | 23/118 | 0.011 |
| 05418 | Fluid shear stress and atherosclerosis | 38/125 | 0.013 |
| 05223 | Non-small cell lung cancer | 22/64 | 0.014 |
| 05034 | Alcoholism | 25/104 | 0.014 |
| 04744 | Phototransduction | 4/9 | 0.015 |
| 05215 | Prostate cancer* | 28/88 | 0.016 |
| 04137 | Mitophagy - animal | 22/61 | 0.018 |
| 04659 | Th17 cell differentiation | 18/72 | 0.024 |
| 03030 | DNA replication* | 13/35 | 0.026 |
| 05031 | Amphetamine addiction | 13/49 | 0.028 |
| 05200 | Pathways in cancer* | 110/434 | 0.031 |
| 04710 | Circadian rhythm | 12/29 | 0.033 |
| 01210 | 2-Oxocarboxylic acid metabolism* | 7/16 | 0.039 |
| 04140 | Autophagy - animal | 36/123 | 0.041 |
| 04372 | Apelin signaling pathway | 34/121 | 0.045 |
| 05226 | Gastric cancer | 36/123 | 0.048 |
| 00515 | Mannose type O-glycan biosynthesis* | 8/20 | 0.048 |
| 03320 | PPAR signaling pathway | 20/56 | 0.048 |

\*Over-representation only

Interestingly, these prednisolone-targeted pathways are closely associated with important skeletal muscle processes such as metabolism, atrophy and regulatory function, alongside previous associations with SMA-related pathways such as FoxO signalling ^66^, p53 signalling ^67^, AMPK signalling ^68^, mitophagy ^69^, circadian rhythm ^70^, PPAR signalling ^71^ and autophagy ^72^ (Table 1). An additional GO analysis also revealed similar skeletal muscle biological processes associated with the DEGs in prednisolone-treated *Smn^-/-^;SMN2* SMA mice such as myotube differentiation, fatty acid oxidation, protein ubiquitination, sarcomere regulation, gluconeogenesis, and circadian rhythm (Table 2; Tables S6-8).

**Table 2.**
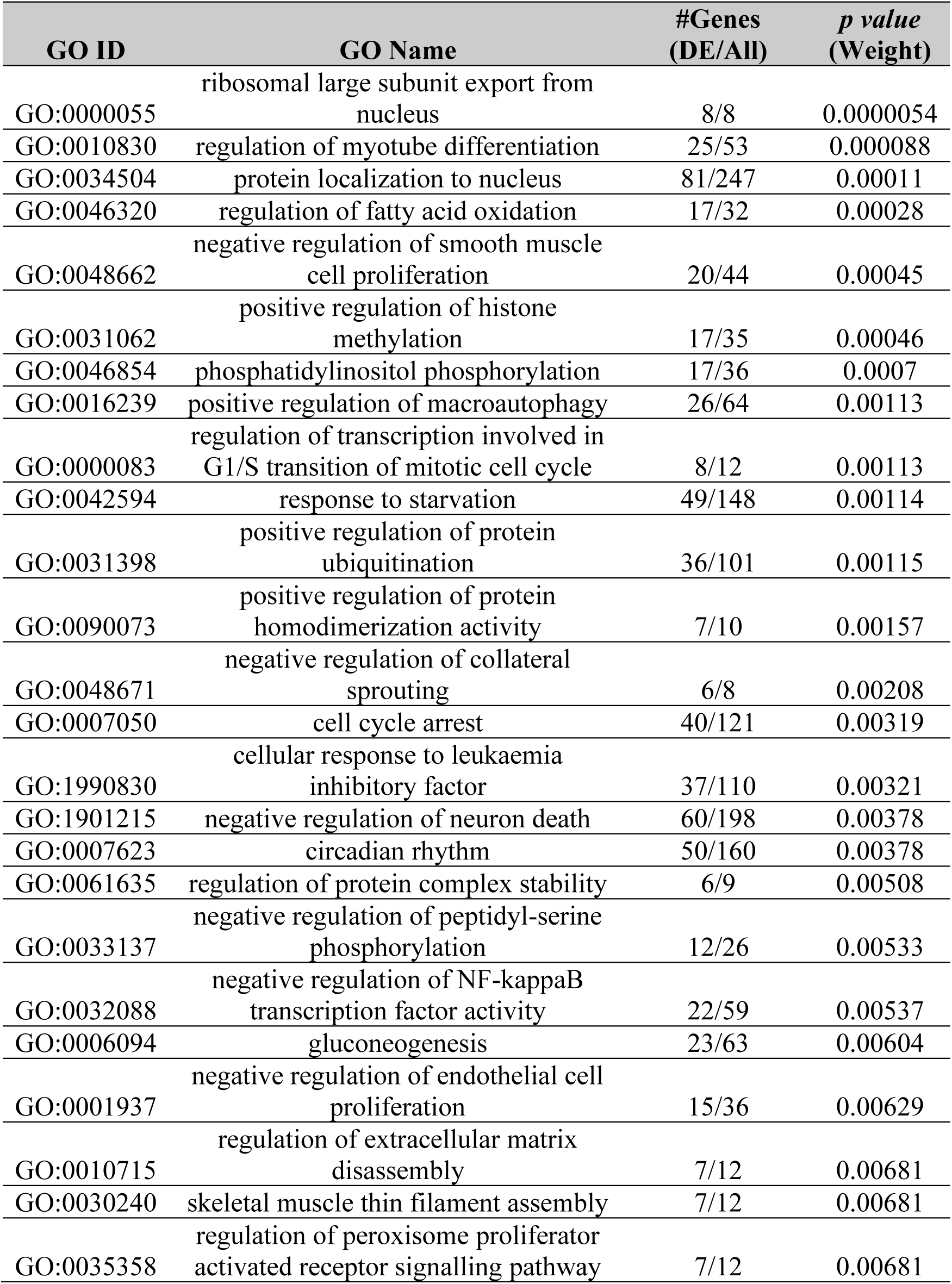
Top Gene Ontology Biological Process pathways targeted in the skeletal muscle of symptomatic prednisolone-treated *Smn^-/-^;SMN2* SMA mice compared with untreated *Smn^-/-^ ;SMN2* SMA mice.

| GO ID | GO Name | #Genes<br>(DE/All) | p value<br>(Weight) |
| --- | --- | --- | --- |
| GO:0000055 | ribosomal large subunit export from nucleus | 8/8 | 0.0000054 |
| GO:0010830 | regulation of myotube differentiation | 25/53 | 0.000088 |
| GO:0034504 | protein localization to nucleus | 81/247 | 0.00011 |
| GO:0046320 | regulation of fatty acid oxidation | 17/32 | 0.00028 |
| GO:0048662 | negative regulation of smooth muscle cell proliferation | 20/44 | 0.00045 |
| GO:0031062 | positive regulation of histone methylation | 17/35 | 0.00046 |
| GO:0046854 | phosphatidylinositol phosphorylation | 17/36 | 0.0007 |
| GO:0016239 | positive regulation of macroautophagy | 26/64 | 0.00113 |
| GO:0000083 | regulation of transcription involved in G1/S transition of mitotic cell cycle | 8/12 | 0.00113 |
| GO:0042594 | response to starvation | 49/148 | 0.00114 |
| GO:0031398 | positive regulation of protein ubiquitination | 36/101 | 0.00115 |
| GO:0090073 | positive regulation of protein homodimerization activity | 7/10 | 0.00157 |
| GO:0048671 | negative regulation of collateral sprouting | 6/8 | 0.00208 |
| GO:0007050 | cell cycle arrest | 40/121 | 0.00319 |
| GO:1990830 | cellular response to leukaemia inhibitory factor | 37/110 | 0.00321 |
| GO:1901215 | negative regulation of neuron death | 60/198 | 0.00378 |
| GO:0007623 | circadian rhythm | 50/160 | 0.00378 |
| GO:0061635 | regulation of protein complex stability | 6/9 | 0.00508 |
| GO:0033137 | negative regulation of peptidyl-serine phosphorylation | 12/26 | 0.00533 |
| GO:0032088 | negative regulation of NF-kappaB transcription factor activity | 22/59 | 0.00537 |
| GO:0006094 | gluconeogenesis | 23/63 | 0.00604 |
| GO:0001937 | negative regulation of endothelial cell proliferation | 15/36 | 0.00629 |
| GO:0010715 | regulation of extracellular matrix disassembly | 7/12 | 0.00681 |
| GO:0030240 | skeletal muscle thin filament assembly | 7/12 | 0.00681 |
| GO:0035358 | regulation of peroxisome proliferator activated receptor signalling pathway | 7/12 | 0.00681 |

Combined, our transcriptomics and pathway analyses suggest that prednisolone treatment attenuated muscle pathologies in SMA mice ^27^ by targeting key muscle metabolism, atrophy and regulatory pathways.

### Drug repositioning algorithms identify novel pharmacological compounds predicted to emulate prednisolone’s activity in skeletal muscle of SMA mice

As mentioned above, while prednisolone treatment significantly improves muscle health and overall disease progression in SMA mice, chronic use of prednisolone can negatively impact skeletal muscle ^35, 39^. As such, we used the DEGs and associated KEGG pathways identified in prednisolone-treated *Smn^-/-^;SMN2* SMA mice to discover alternative drugs predicted to mimic prednisolone’s molecular effects in SMA skeletal muscle. Initially, we utilized the in-built integration of KEGG drugs database in iPathwayGuide ^53^ and the DGIdb v3.0 ^58^ database to initially reveal a total of 580 compounds (Tables S9-12). To filter down our list, we focused on the drug compounds 1) that targeted > 5 prednisolone-targeted pathways or linked to upstream regulators, 2) were clinically approved and 3) were not associated with promotion of muscle-wasting (e.g., primary anti-cancer drugs ^73^), leaving a total of 20 potential candidates (Tables 3-4). Interestingly, our combined *in silico* drug repositioning approach revealed a subset of candidates previously investigated in SMA such as celecoxib ^74^ (ClinicalTrials.gov ID: NCT02876094) and colforsin ^75^. To further validate our bioinformatics strategy, we chose to continue our study with drugs not yet assessed for SMA, focusing on those previously used safely in young patients and orally bioavailable. With these criteria, we narrowed down our selection to metformin, a generic asymmetric dimethyl-biguanide type 2 diabetes mellitus (T2DM) drug ^76^ and oxandrolone, a synthetic anabolic steroid with a higher ratio of anabolic: androgynous effects for further study ^77^. Thus, using a transcriptomics-based *in silico* drug repositioning platform, we were able to generate a list of clinically approved pharmacological compounds that are predicted to emulate prednisolone’s activity in skeletal muscle.

**Table 3.** Top 10 clinically approved drugs identified by KEGG database based on prednisolone-targeted KEGG pathways in symptomatic prednisolone-treated *Smn^-/-^;SMN2* SMA mice.

| Pathways Targeted | KEGG ID | Compound | Bioavailability | Gene Targets |
| --- | --- | --- | --- | --- |
| 10 | D03297 | Mecasermin<br>(genetical recombination)<br>(JAN) | Subcutaneous | *Igf1r* |
| 10 | D04870 | Mecasermin<br>rinfabate<br>(USAN/INN) | Intravenous | *Igf1r* |
| 9 | D09680 | Teprotumumab<br>(USAN/INN) | Intravenous | *Igf1r* |
| 8 | D04966 | Metformin<br>(USAN/INN) | Oral | *Prkag3* |
| 8 | D00944 | Metformin<br>hydrochloride<br>(JP16/USP) | Oral | *Prkag3* |
| 7 | D01697 | Colforsin<br>daropate<br>hydrochloride<br>(JAN) | Oral | *Adcy1*,<br>*Adcy6*<br>*Adcy7* |
| 6 | D01146 | Iguratimod<br>(JAN/INN) | Oral | *Nfkb1* |
| 5 | D07058 | Acamprosate<br>(INN) | Oral | *Grin1*,<br>*Grin2a*,<br>*Grin2b*,<br>*Grin2c*,<br>*Grin2d* |
| 5 | D02754 | Acitretin<br>(USP/INN);<br>Soriatane (TN) | Oral | *Rarb*,<br>*Rxrb*,<br>*Rxrg* |
| 5 | D00085 | Insulin<br>(JAN/USP) | Intravenous<br>Subcutaneous | *Insr* |

**Table 4.** Top 10 clinically approved drugs identified by DGIdb database based on prednisolone-targeted KEGG pathways in symptomatic prednisolone-treated *Smn^-/-^;SMN2* SMA mice.

| Compound | Bioavailability | Gene Target |
| --- | --- | --- |
| Testosterone | Busal<br>Nasal<br>Oral<br>Topical<br>Transdermal<br>Subcutaneous | *Ar* |
| Oxandrolone | Oral | *Ar* |
| Nandrolone<br>phenpropionate | Oral | *Ar* |
| Progesterone | Oral<br>Vaginal<br>Intramuscular | *Ers1* |
| Tibolone | Oral | *Ers1* |
| Cannabidiol | Oral<br>Inhale | *Cnr1* |
| Insulin, neutral | Intravenous<br>Intramuscular<br>Subcutaneous | *Insr* |
| Celecoxib | Oral | *Pdpk1* |
| Tocilizumab | Intravenous<br>Subcutaneous | *Il-6ra* |
| Sarilumab | Subcutaneous | *Il-6ra* |

### Metformin’s primary predicted target gene *Prkag3* is dysregulated in skeletal muscle of both severe *Smn^-/-^;SMN2* and intermediate *Smn^2B/-^* SMA mice

As previously mentioned, metformin is an orally administered T2DM drug that we selected as one of the candidates to validate our bioinformatics-based drug repositioning approach. Importantly, metformin has over 60 years of clinical use with a well-known safety profile ^76^ and recorded administration in younger patients ^78^. Furthermore, it has been previously repositioned and conferred ergogenic activities in muscular disorders such as DMD ^79^ and congenital muscular dystrophy type 1 A (CMDT1A) ^80^, highlighting its potential as a skeletal muscle therapy.

Our iPathwayGuide analysis predicted that metformin could emulate prednisolone’s targeting of the KEGG: 04068 FoxO signalling pathway (Figure S2.a). In particular, metformin was predicted to mimic prednisolone’s upregulation of *Prkag3*, which encodes for the AMPK-γ3 subunit of the predominant skeletal muscle AMPK-α2β2γ3 isoform complex ^81^ (Figure S2). Furthermore, *Prkag3* upregulation was predicted to coherently downregulate the expression of *FoxO1*, *FoxO3* and *Foxo4* isoforms, whilst upregulating *FoxO6* (Figure S2) supporting previous literature associating these FoxO isoforms with promotion of muscle atrophy ^66, 82^. Importantly, the expression pattern of these genes in the prednisolone-treated *Smn^-/-^;SMN2* SMA mice were normalized to healthy *Smn^+/-^;SMN2* levels (Figure S2.c), supporting the usefulness of investigating metformin and these targets in SMA skeletal muscle.

We thus measured the mRNA expression levels of *Prkag3* and *FoxO* isoforms in the triceps of both symptomatic P7 severe *Smn^-/-^;SMN2* and P19 intermediate *Smn^2B/-^* SMA mice alongside their respective healthy controls. We indeed observed that *Prkag3* levels were significantly downregulated in skeletal muscle of both SMA mouse models (Figure 2.a), supporting the bioinformatics data. However, none of the *FoxO* isoforms were significantly different between SMA mice and their healthy controls (Figures 2.b-e). Previous research also reported no significant upregulation of *FoxO* isoforms in P7 severe *Smn^-/-^;SMN2* SMA mice via qPCR ^66^. However, the fact that our qPCR data did not reflect the bioinformatics predictions may be due to variability in our experimental cohorts, the sequencing depth coverage not being sufficiently conservative and/or intrinsic differences between RNA-Seq and primer-based qPCR approaches ^83^.

**Figure 2.**
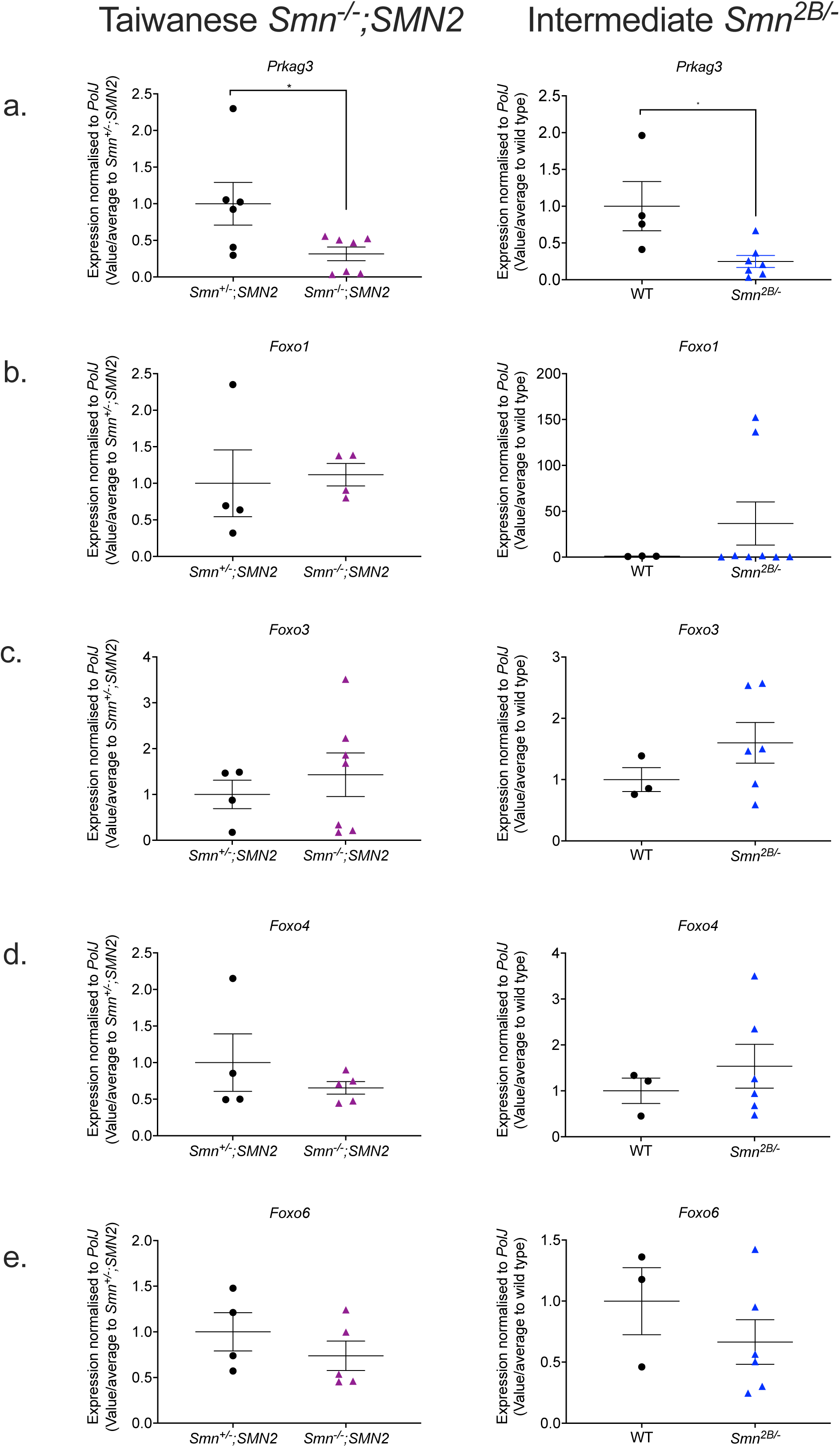
The metformin target gene, *Prkag3,* is significantly downregulated in the skeletal muscle of both symptomatic severe *Smn^-/-^;SMN2* and intermediate *Smn^2B/-^* SMA mice. qPCR analysis of mRNA levels for predicted metformin target genes a. *Prkag3*, b. *Foxo1*, c. *Foxo3*, d. *Foxo4* and e. *Foxo6* in the harvested triceps of symptomatic untreated P7 Taiwanese *Smn^-/-^;SMN2* SMA mice (violet) and healthy *Smn^+/-^;SMN2* controls (black) (left panel) and symptomatic untreated P19 intermediate *Smn^2B/-^* SMA mice (blue) and wild type (C57BL/6J background) controls (black) (right panel). Data are shown as scatter plot represent as mean ± SEM error bars; n = 4-7 animals per experimental group, unpaired t-test, \**p* <0.05. *Smn^-/-^;SMN2 Prkag3*: *p* = 0.04; *Smn^-/-^;SMN2 Foxo1*: *p* = 0.82; *Smn^-/-^;SMN2 Foxo3*: *p* = 0.54; *Smn^-/-^;SMN2 Foxo4*: *p* = 0.37; *Smn^-/-^;SMN2 Foxo6*: *p* = 0.34; *Smn^2B/-^ Prkag3*: *p* = 0.02; *Smn^2B/-^ Foxo1*: *p* = 0.39; *Smn^2B/-^ Foxo3*: *p* = 0.27; *Smn^2B/-^ Foxo4*: p = 0.48; *Smn^2B/-^ Foxo6*: *p* = 0.33.

Overall, our qPCR experiments revealed that the primary metformin target *Prkag3* matched its bioinformatics prediction of being downregulated in both severe *Smn^-/-^;SMN2* and intermediate *Smn^2B/-^* SMA mice, suggesting that this gene may be involved in both severe and milder SMA pathologies and an appropriate therapeutic molecular target in SMA muscle.

### The predicted target genes for metformin are mostly Smn-independent in an SMA muscle cellular model

We next wanted to better understand if the aberrant expression of the metformin target genes was dependent on SMN expression and/or muscle atrophy. Thus, we firstly generated siRNA-mediated Smn-depleted C2C12 myoblast-like cells, a useful and previously successful *in vitro* model ^84^. We confirmed by qPCR that *Smn* mRNA levels were significantly reduced by up to 90% in C2C12 myoblasts and D8 C2C12 myotubes compared to scrambled siRNA and untreated controls (Figure S3). We next investigated the effects of Smn knockdown on the expression of the predicted metformin target genes. In C2C12 myoblasts, we identified a significant upregulation of only the *FoxO3* gene in Smn-depleted C2C12 myoblasts compared to controls (Figure 3.a), which reflects previous microarray analyses of specific *FoxO* isoforms upregulated in quadriceps femoralis muscle biopsies from type 1 SMA patients ^85^. However, in C2C12 myotubes we found that Smn knockdown (KD) had no effect on the expression of predicted metformin target genes (Figure 3.b), suggesting that for the most part, the expression of the predicted metformin genes is Smn- independent, thus representing ideal targets for SMN-independent therapies.

**Figure 3.**
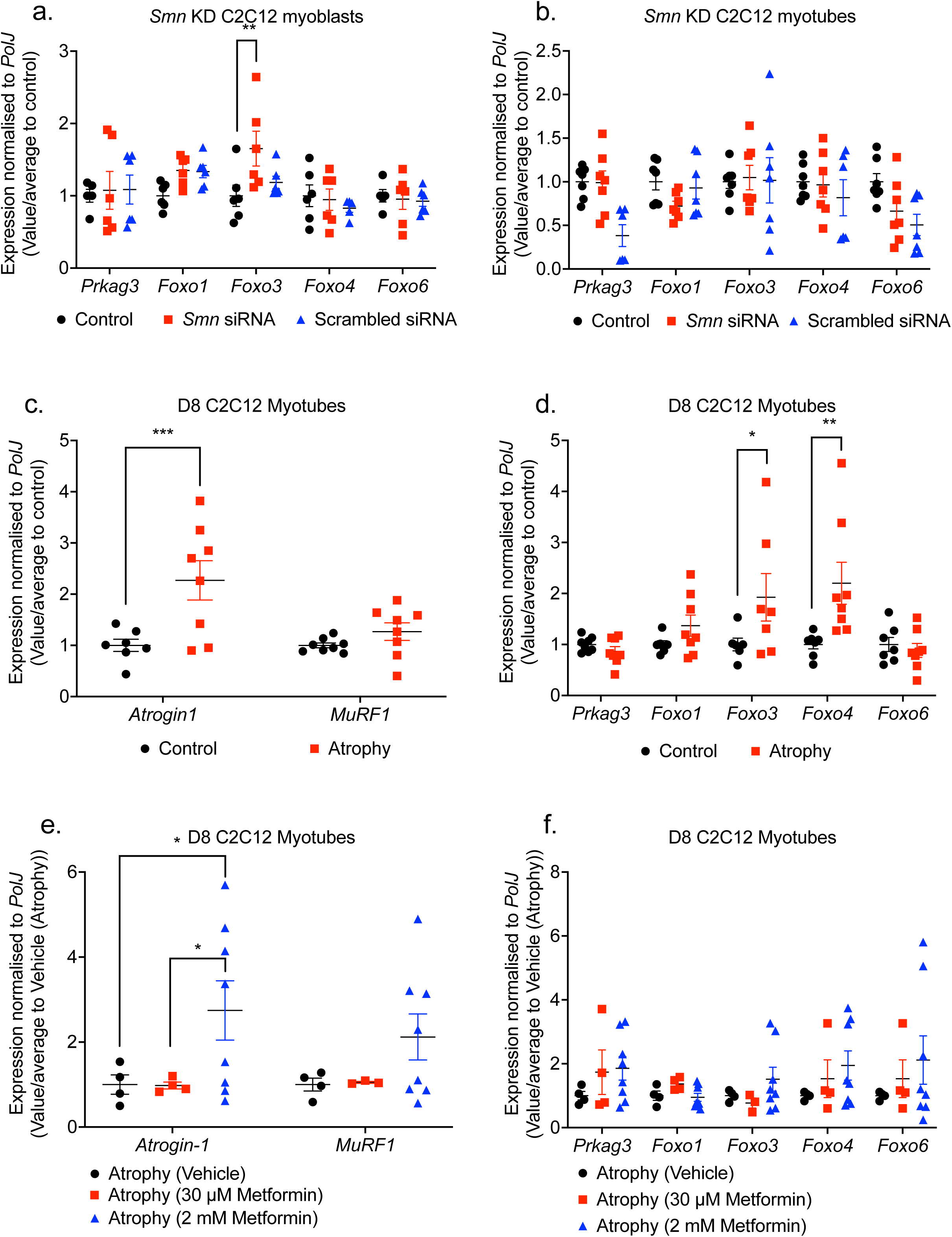
Metformin target genes are pre-dominantly SMN-independent in an SMA muscle C2C12 cellular model. *Smn* siRNA knockdown (red) was performed for **a.** 48 hours in C2C12 myoblasts and **b.** every 48 hours throughout differentiation in D8 C2C12 myotubes. mRNA expression of metformin target genes *Prkag3*, *Foxo1*, *Foxo3*, *Foxo4* and *Foxo6* was measured by qPCR and compared to non- transfected (black) and scrambled siRNA transfected controls (blue). D8 C2C12 myotubes were serum-starved for 24 hours to induce canonical atrophy (red). mRNA expression of **c.** atrogenes *Atrogin-1* and *MuRF-1* and **d.** Metformin target genes *Prkag3*, *Foxo1*, *Foxo3*, *Foxo4* and *Foxo6* was measured by qPCR and compared against non-starved myotubes (black). Serum-starved D8 C2C12 myotubes were treated with either physiological 30 μM metformin (red) or supraphysiological 2 mM metformin (blue) for 24 hours to evaluate mRNA expression via qPCR of **e.** atrogenes *Atrogin-1* and *MuRF-1* and **f.** Metformin target genes *Prkag3*, *Foxo1*, *Foxo3*, *Foxo4* and *Foxo6* compared to serum-starved PBS vehicle treated control (black). Data are shown as scatter plot that represent mean ± SEM error bars; n = 4 samples per group across two independent experiments. Two-way ANOVA followed by uncorrected Fisher’s least significant difference (LSD). F = 3.543 (**a.**); F = 2.332 (**b.**); F = 4.9 (**c.**); F = 3.493 (**d.**); F = 0.057 (**e.**); F = 0.235 (**f.**); F = 0.401, \**p* < 0.05, \*\**p* < 0.01, \*\*\**p* < 0.001.

We next investigated if the expression of the predicted metformin target genes is affected *in vitro* by muscle atrophy. However, one difficulty in mimicking SMA muscle atrophy *in vitro* is establishing denervation. Thus, based on evidence of shared pathway similarities from different pro-atrophy factors such as starvation and denervation ^86^, we used a validated method of 24-hour serum-starvation in C2C12 myotubes to induce canonical atrophy, as confirmed by myotube loss and upregulation of pro-atrophic *atrogin-1* levels (Figure 3.c). Next, we evaluated the expression of the predicted metformin target genes and observed a significant upregulation of *FoxO3* and *FoxO4* isoforms (Figure 3.d), reflecting their established roles in atrophy-dependent ubiquitin- proteasome pathways ^66^.

We then evaluated whether metformin could attenuate muscle atrophy in C2C12 myotubes. Based on initial gene-dose response experiments in both control C2C12 myoblasts and D8 myotubes, we treated our cells with physiological (60 μM) and supraphysiological (2 mM) metformin concentrations for 24 hours (Figures S4) ^88^. The 30 μM physiological metformin concentration for 24 hours did not attenuate muscle atrophy or impact the expression of the target genes in the serum starved C2C12 myotubes (Figure 3.e-f). However, for the supraphysiological 2 mM metformin concentration ^88^, we observed an upregulation of *Atrogin-1* levels (Figure 3.e), suggesting an exacerbation of muscle atrophy. Further analysis of the predicted metformin target genes revealed no significant impact on their expression patterns either (Figure 3.f), suggesting that exacerbation of atrophy in C2C12 myotubes by supraphysiological metformin concentrations involve factors mostly outside of our predicted targets. However, it should be noted that metformin may have different effects in SMA muscle as there are still differences between distinct pro-atrophic factors^86^.

Overall, our *in vitro* studies revealed that although most of our predicted metformin target genes are SMN-independent with some linked to muscle atrophy, they were mostly not linked to metformin’s influence on canonical atrophy in C2C12 myotubes.

### Dose-dependent effect of metformin on disease progression and survival in *Smn^2B/-^*SMA mice

Next, we assessed metformin in the intermediate *Smn^2B/-^* SMA mouse model ^43^. The rationale for conducting our *in vivo* pharmacological studies in the *Smn^2B/-^* SMA mice was based on their longer lifespan ^43^, responsiveness to SMN-independent therapies ^42, 70^, established metabolic and myopathy defects ^43, 66, 70, 89–91^, and later symptomatic onset ^43^ making them a clinically relevant model for starting treatment regimens >P5 time-points ^42, 70^.

We initially administered a 200 mg/kg daily dose (diluted in 0.9% saline) starting from P5 until humane endpoint in *Smn^2B/-^* SMA mice and *Smn^2B/+^* healthy control littermates, based on this concentration’s previous success in the muscle disorder CMDT1A ^80^. We observed no significant improvement in survival of *Smn^2B/-^* SMA mice treated with 200 mg/kg/day compared to untreated (Figure 4.a) and vehicle-treated animals (Figure S5). We also found a significant reduction in the body weight of 200 mg/kg/day metformin-treated *Smn^2B/-^* SMA mice compared to untreated SMA animals, beginning 4 days after initial treatment at P9 (Figure 4.b.) However, we did not observe any effects on weight in the 200 mg/kg/day metformin treated *Smn^2B/+^* healthy control mice (Figure S6.b), indicating a disease specific effect of metformin. In terms of motor function, there was no significant difference in the righting reflex between untreated and 200 mg/kg/day metformin-treated *Smn^2B/-^* SMA mice (Figure 4.c) and *Smn^2B/+^* healthy control animals (Figure S6.a-c).

**Figure 4.**
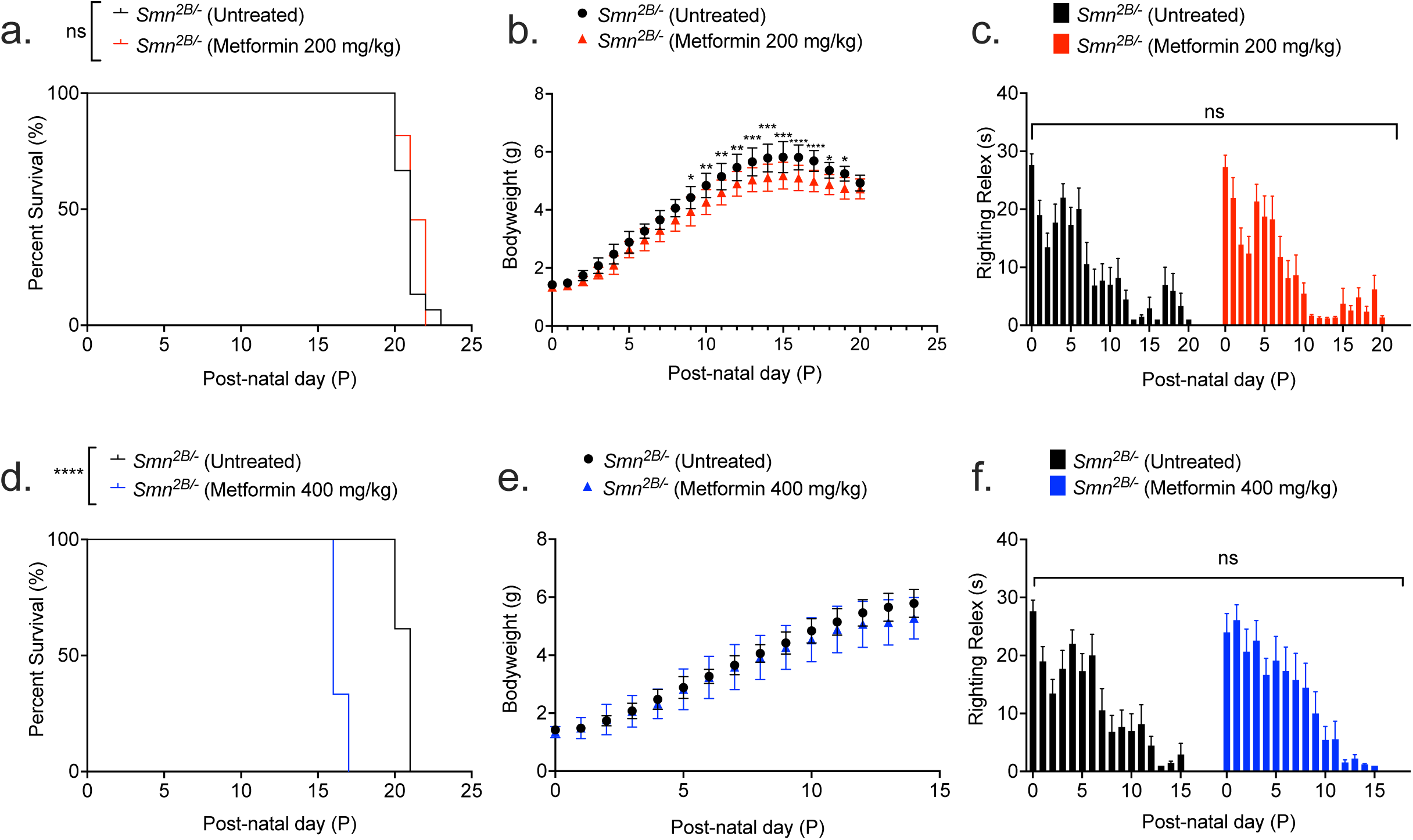
200 mg/kg/day metformin does not improve disease phenotype, while 400 mg/kg/day metformin reduces survival. All treated animals received a daily dose of metformin (either 200 or 400 mg/kg/day, diluted in 0.9% saline) by gavage starting at P5. **a.** Survival curves of untreated (n = 13, 21 days median survival, black) and 200 mg/kg/day metformin-treated (n = 11, 21 days median survival, red) *Smn^2B/-^* SMA mice. Kaplan-Meier survival curve shown with log rank (Mantel-Cox) test, ns = not significant, *p* = 0.237. **b.** Daily weights of untreated (n = 13, black) and 200 mg/kg/day metformin- treated (n = 11, red) *Smn^2B/-^* SMA mice. Data represented as mean ± SEM error bars; Two-way ANOVA followed by a Sidak’s multiple comparison test, F = 402.1, df = 455, \**p* < 0.05, \*\**p* < 0.01, \*\*\**p* < 0.001, \*\*\*\**p* < 0.0001. **c.** Daily righting reflex test for motor function activity up to a 30 second maximum time point in untreated (n = 13, black) and 200 mg/kg/day metformin- treated (n = 11, red) *Smn^2B/-^* SMA mice. Data are shown as bar chart with mean ± SEM error bars; unpaired T-test, ns = not significant, *p* = 0.833**. d.** Survival curves of untreated (n = 13, 21 days median survival, black) and 400 mg/kg/day metformin-treated (n = 4, 16 days median survival, blue) *Smn^2B/-^* SMA mice. Kaplan-Meier survival curve shown with log rank (Mantel-Cox) test, \*\*\*\**p* < 0.0001. **e.** Daily weights of untreated (n = 13, black) and 400 mg/kg/day metformin- treated (n = 9, blue) *Smn^2B/-^* SMA mice. Data represented as mean ± SEM error bars; Two-way ANOVA followed by a Sidak’s multiple comparison test, F = 184.9.1, df = 300. **f.** Daily righting reflex test for motor function activity up to a 30 second maximum time point in untreated (n = 13, black) and 400 mg/kg/day metformin-treated (n = 9, blue) *Smn^2B/-^* SMA mice. Data are shown as bar chart with mean ± SEM error bars; unpaired T-test, ns = not significant, *p* = 0.733.

Since our initial 200 mg/kg/day metformin dosage did not improve disease onset or disease progression in *Smn^2B/-^* SMA mice, we conducted pilot studies with a later treatment start point (P8) and a lower dose (100 mg/kg/day), which demonstrated similar effects to our initial dosing regimen (data not shown). We therefore tried a higher daily dosage of 400 mg/kg/day, starting at P5. Surprisingly, the higher 400 mg/kg/day dose significantly reduced survival in *Smn^2B/-^* SMA pups by 5 days (Figure 4.d), while having no significant impact on weight or righting reflex (Figures 4.e-f). Interestingly, 400 mg/kg/day metformin had no adverse effects in the healthy *Smn^2B/+^*control littermates (Figures S6.d-f), suggesting that the adverse effects of the higher dose of metformin is disease specific.

Thus, our *in vivo* experiments demonstrated that metformin did not emulate prednisolone’s beneficial effects on SMA disease progression in SMA mice and is in fact not an ideal therapy candidate, due to dose- and disease-dependent adverse effects in SMA.

### Higher 400 mg/kg/day metformin dosage is associated with hypoglycaemia in non-fasted *Smn^2B/-^* SMA mice

We next investigated the potential causes behind metformin’s adverse effects in SMA mice. With metformin being a glucose lowering agent for T2DM, we initially assessed blood glucose levels in P14 non-fasted, untreated, 200- and 400-mg/kg/day metformin treated *Smn^2B/-^* SMA and *Smn^2B/+^* healthy mice 2 hours after the final treatment. This time point was chosen to account for the reduced median survival in the higher dose SMA cohort. We observed that neither 200- or 400- mg/kg/day metformin treatments lowered blood glucose levels in *Smn^2B/+^* healthy mice (Figure 5.a). However, we reported a significant reduction in blood glucose levels in 400 mg/kg/day metformin-treated *Smn^2B/-^* SMA mice compared to untreated SMA animals (Figure 5.a). Our results suggest that hypoglycaemic shock could have been one of the possible causes behind the premature death in the 400 mg/kg/day SMA cohort, further exacerbating the previously reported hypoglycaemia in SMA models ^89^ and patients ^92, 93^.

**Figure 5.**
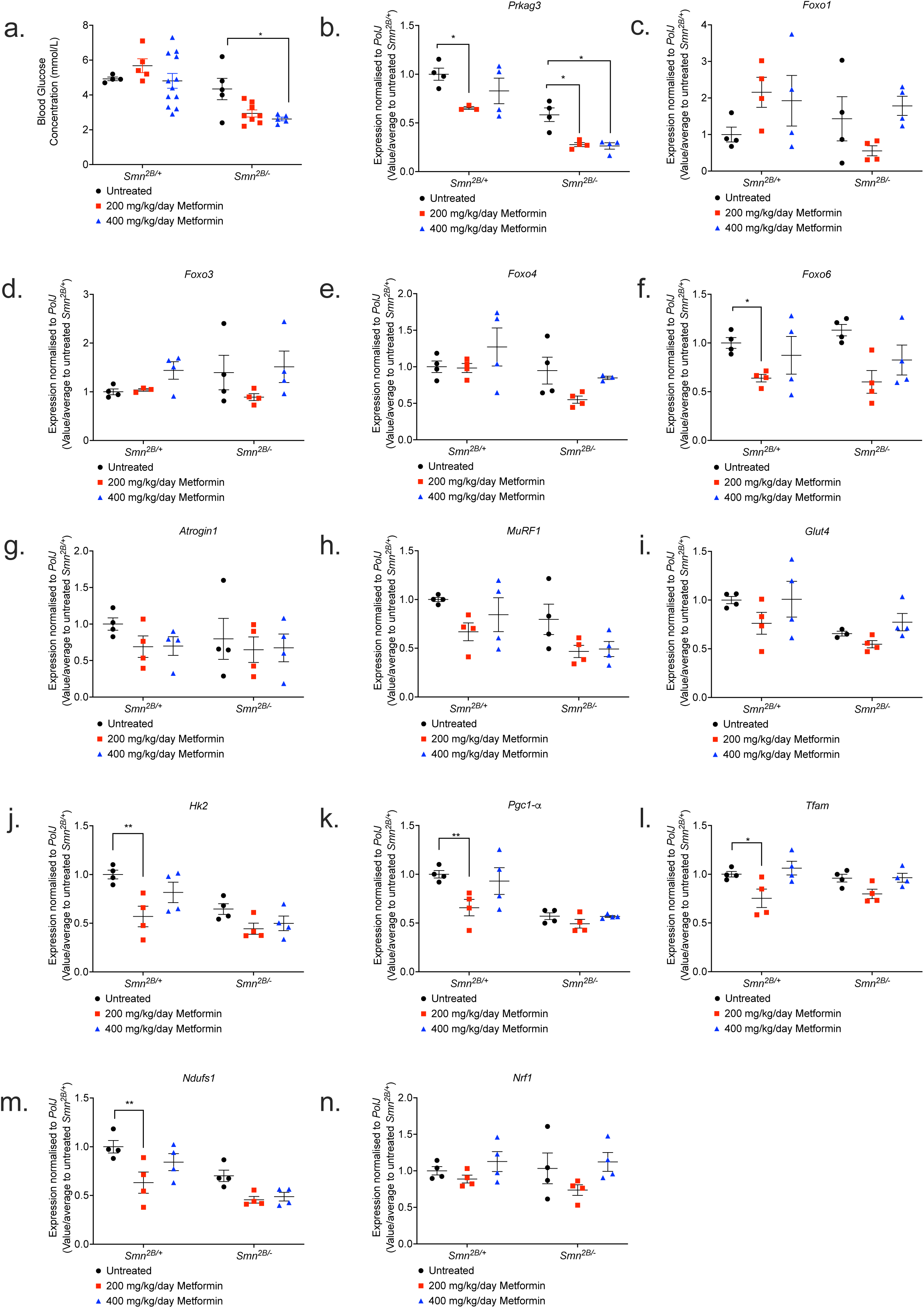
400 mg/kg/day metformin significantly lowers blood-glucose levels in *Smn^2B/-^* SMA mice, with no impact on markers of atrophy, glucose metabolism and mitochondrial regulation in skeletal muscle. **a.** Blood-glucose concentrations (mmol/L) were measured 2 hours after final treatment from untreated (black) and 200 (red) or 400 mg/kg/day metformin-treated (blue), non-fasted P14 *Smn^2B/+^*healthy and *Smn^2B/-^* SMA mice. Data represented as bar chart with scatter graph represented as mean ± SEM error bars, n = 4 animals per group; two-way ANOVA with Tukey’s multiple comparisons test, F = 25.49, * *p* <0.05. qPCR analysis of predicted metformin target genes b. *Prkag3*, c. *Foxo1*, **d.** *Foxo3*, e. *Foxo4*, f. *Foxo6*; atrogenes **g.** *Atrogin-1*, h. *MuRF1*; and glucose uptake and metabolism genes i. *Glut4*, j. *Hk2*; and mitochondrial regulatory genes k. *Pgc1-α,* l. *Tfam,* m. *Ndufs1* and n. *Nrf1* in the TA muscle from untreated and 200 or 400 mg/kg/day metformin-treated, P14 *Smn^2B/+^* healthy and *Smn^2B/-^* SMA mice. Data are shown as scatter graph represented as mean ± SEM error bars, n = 4 animals per group; two way ANOVA with Tukey’s multiple comparisons test, *Prkag3* (F = 59.92), *Foxo1* (F = 1.507), *Foxo3* (F = 0.343), *Foxo4* (F = 6.475), *Foxo6* (F = 0.024), *Atrogin-1* (F = 0.381), *MuRF1* (F = 7.838), *Glut4* (F = 9.9), *Hk2* (F = 17.78), *Pgc1-α* (F = 29.84), *Tfam* (F = 0.423) *Ndufs1* (F = 22.66), and *Nrf1* (F = 0.164).

### Metformin did impact the expression of the predicted target genes, but not muscle pathology markers in *Smn^2B/-^* SMA mice

We next evaluated the effect of metformin on the expression of the predicted target genes in TAs from P14 untreated, 200- and 400-mg/kg/day metformin-treated *Smn^2B/-^* SMA and *Smn^2B/+^* healthy mice 2 hours after final treatment. Contrary to the drug repositioning prediction that metformin could reverse *Prkag3* downregulation in SMA muscle (Figure S3), we instead discovered that both 200- and 400-mg/kg/day metformin doses exacerbated *Prkag3* downregulation in *Smn^2B/-^* SMA muscle (Figure 5.b). Furthermore, although metformin had no impact on *FoxO1*, *FoxO3* and *FoxO4* isoforms in both *Smn^2B/+^* healthy and *Smn^2B/-^* SMA muscle (Figures 5.c-e), for the 200 mg/kg/day *Smn^2B/-^* SMA cohort, metformin significantly reduced *FoxO6* expression (Figure 5.f), which again contrasted our bioinformatics prediction that metformin would upregulate the expression of this isoform in SMA muscle. Thus, the drug-gene response for metformin-treated *Smn^2B/-^* SMA skeletal muscle reveals a contrasting pattern that does not match the bioinformatic predictions.

We next investigated metformin’s effects on the expression of dysregulated molecular markers associated with muscle atrophy (*Atrogin-1* and *MuRF-1*) and glucose metabolism (*Glut4* and *Hk2*) ^94, 95^. We however observed that neither atrophy (Figures 5.g-h) or glucose metabolism markers (Figures 5.i-j) were affected by 200- and 400-mg/kg/day metformin treatments in the *Smn^2B/-^* SMA mice when compared to untreated animals.

We also investigated markers associated with mitochondrial biogenesis and function in muscle (*Pgc1-α, Tfam*, *Ndufs1*, and *Nrf1*), as previous research has established that these features are impaired in SMA skeletal muscle ^69^ and a common mechanism of action for metformin is mild inhibition of mitochondrial electron transport complex 1 (or NADH:ubiquinone oxidoreductase)^96^. We found that that neither the 200- or 400-mg/kg/day dose of metformin impacted the expression of mitochondrial genes in the skeletal muscle of *Smn^2B/-^* SMA mice (Figures 5.k-n), although we do observe a significant downregulation of *Pgc1-α, Tfam*, and *Ndufs1* in the skeletal muscle of 200 mg/kg/day metformin-treated *Smn^2B/+^* healthy counterparts (Figures 5.k-n).

Overall, our data highlights that metformin did not have a direct impact on the predicted target genes in skeletal muscle of SMA mice. Furthermore, the absence of direct impact on muscle atrophy, glucose metabolism, and mitochondrial function markers following metformin treatment in *Smn^2B/-^* SMA muscle, suggests that the adverse effects associated with the 400 mg/kg/day dosage may not have been linked to muscle-intrinsic effects.

### A higher dose of metformin is associated with dysregulation of mitochondrial regulatory genes in the spinal cord of *Smn^2B/-^* SMA mice

We next investigated the effects of metformin on the spinal cord given that metformin is systemically distributed ^97^, has the ability to cross the blood-brain-barrier (BBB) ^98^ and can impact the mitochondria in the spinal cord ^99^. We thus evaluated whether metformin altered the expression of mitochondrial markers (*Pgc1-α*, *Tfam*, *Nrf1* and *Ndufs1*) in the spinal cord of P14 untreated, 200- and 400-mg/kg/day metformin treated *Smn^2B/-^* SMA mice compared to *Smn^2B/+^* healthy mice, 2 hours after final treatment.

For *Pgc1-α*, a master regulator of mitochondrial biogenesis and function, we observed that although both 200- and 400-mg/kg/day metformin concentrations significantly reduced its expression levels in *Smn^2B/+^* healthy spinal cords (Figure 6.a), it was only the higher concentration that significantly reduced *Pgc1-α* expression in *Smn^2B/-^* SMA spinal cords (Figure 6.a). Similarly, 400 mg/kg/day metformin significantly reduced *Ndufs1* levels in both *Smn^2B/+^* healthy and *Smn^2B/-^*SMA spinal cords (Figure 6.b), suggesting that for these mitochondrial health markers the higher metformin dose negatively affected their expression independent of disease status. On the other hand, although not affected by metformin in *Smn^2B/-^* SMA spinal cords, *Nrf1* gene expression was significantly reduced by both metformin doses in the spinal cord of *Smn^2B/+^* healthy mice (Figure 6.c), whilst *Tfam* was not affected by metformin in either cohort (Figure 6.d).

**Figure 6.**
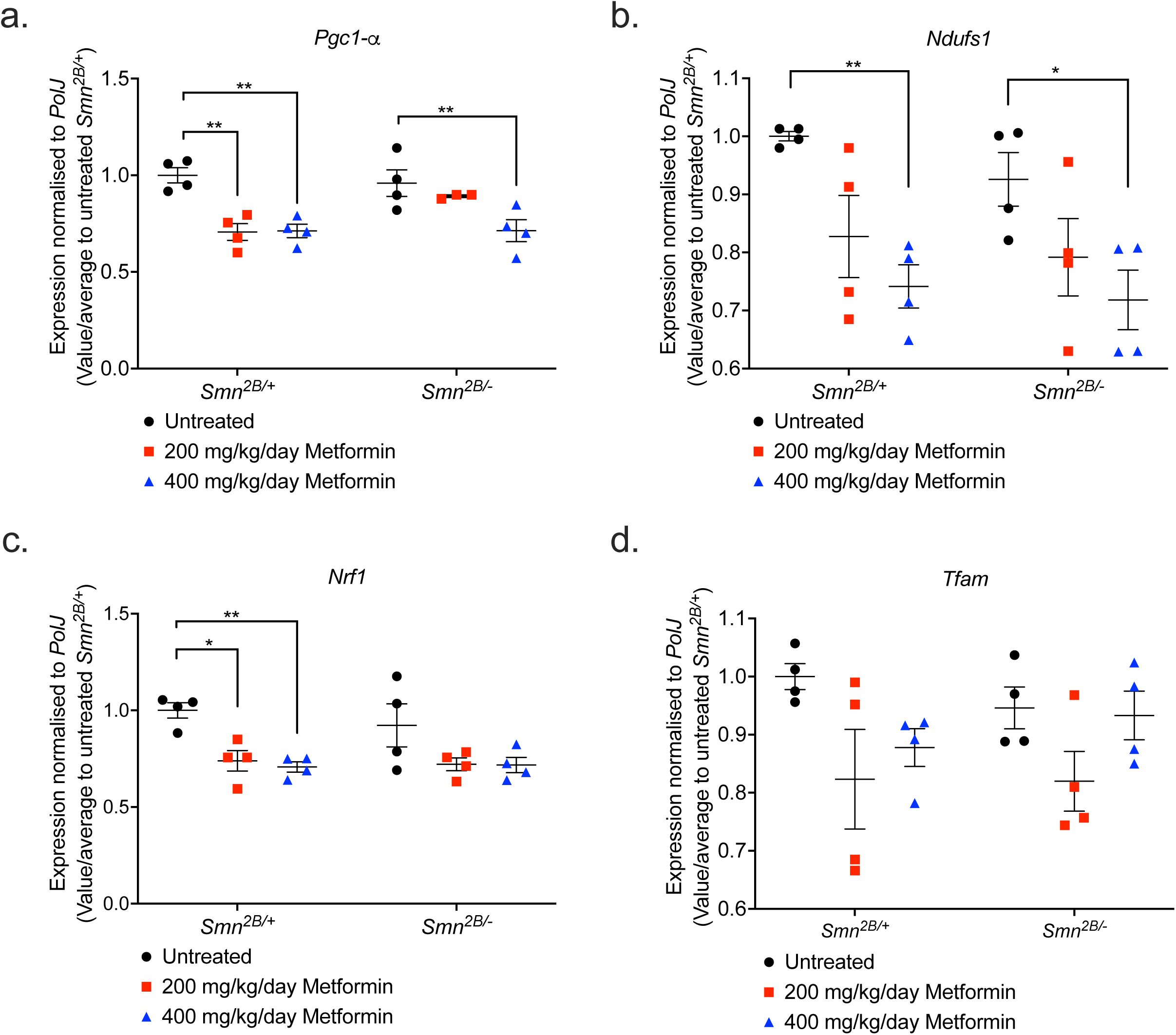
400 mg/kg/day metformin dysregulates mitochondrial regulatory genes exclusively in spinal cord tissue from Smn*^2B/-^* SMA mice. qPCR analysis of mitochondrial regulatory genes a. *Pgc1-α,* **b.** *Ndufs1,* **c.** *Nrf1* and **d.** *Tfam* in the spinal cord from untreated (black) and 200 (red) or 400 mg/kg/day metformin-treated (blue), P14 *Smn^2B/+^* healthy and *Smn^2B/-^* SMA mice. Data are shown as scatter graph represented as mean ± SEM error bars, n = 4 animals per group; two-way ANOVA with Tukey’s multiple comparisons test. a. *Pgc1-α* (F = 1.526), b. *Ndufs1* (F = 1.135), **c.** *Nrf1* (F = 0.362) and d. *Tfam* (F = 0.614), \**p* < 0.05, \*\**p* < 0.01.

Our results demonstrating that the higher dose of metformin (400 mg/kg/day) appears to specifically dysregulate certain mitochondrial genes in the spinal cord of *Smn^2B/-^* SMA mice is supported by recent evidence of tissue-dependent differences in conserved cellular processes between SMA motor neurons and skeletal muscle ^72^. Thus, although further in-depth investigations would be needed, our results on mitochondrial health markers suggest that metformin’s adverse effects in SMA mice could be linked to the exacerbation of neuronal mitochondrial dysfunction.

### Oxandrolone’s predicted target gene, *Ddit4,* is dysregulated in the skeletal muscle of severe *Smn^-/-^;SMN2* SMA mice

Our second drug candidate that we selected to mimic prednisolone activities was oxandrolone, a synthetic orally bioavailable anabolic steroid that confers minimal androgynous effects ^77^. Importantly for SMA, oxandrolone has been successful in the promotion of muscle growth for DMD ^100^ and mixed gender burn injury patients ^101^.

Oxandrolone was predicted to upregulate the *Ar* gene in SMA muscle (Figure S7). The upregulation of *Ar* was predicted to directly upregulate downstream effectors *Igfbp5* and *myogenin* (or *MyoG*) (Figure S7), which both regulate muscle differentiation, regeneration and myofiber growth ^102, 103^. Furthermore, *Ar* was predicted to indirectly upregulate *Dok5*, a signalling protein linked to insulin and IGF-1 activity ^104^ and *Akap6*, which is involved in the modulation of muscle differentiation and regeneration ^105^ (Figure S7). In addition to these factors, we also decided to investigate *Ddit4* as an oxandrolone target based on its direct relation with *Ar* ^106^ and being one of the top 20 downregulated DEG targets of prednisolone in *Smn^-/-^;SMN2* SMA skeletal muscle (Table S4; Figure S7).

Similar to our metformin strategy above, we initially wanted to evaluate the mRNA expression levels of these target genes in the triceps of both symptomatic P7 severe *Smn^-/-^;SMN2* and P19 intermediate *Smn^2B/-^* SMA mice alongside their respective healthy controls. Overall, we identified no significant dysregulated expression of the target genes *Ar*, *Akap6*, *Igfbp5*, *Dok5* and *MyoG* in both severe *Smn^-/-^;SMN2* and intermediate *Smn^2B/-^* SMA mice (Figures 7.a-e). However, for *Ddit4,* we did identify a significant upregulation only in *Smn^-/-^;SMN2* SMA mice (Figures 7.f), supporting both our bioinformatics data for this gene and its known pro-atrophic role ^106^, indicating that it may play an important role in SMA muscle pathologies. In summary, the majority of the predicted oxandrolone target genes did not significantly reflect their bioinformatic predictions.

**Figure 7.**
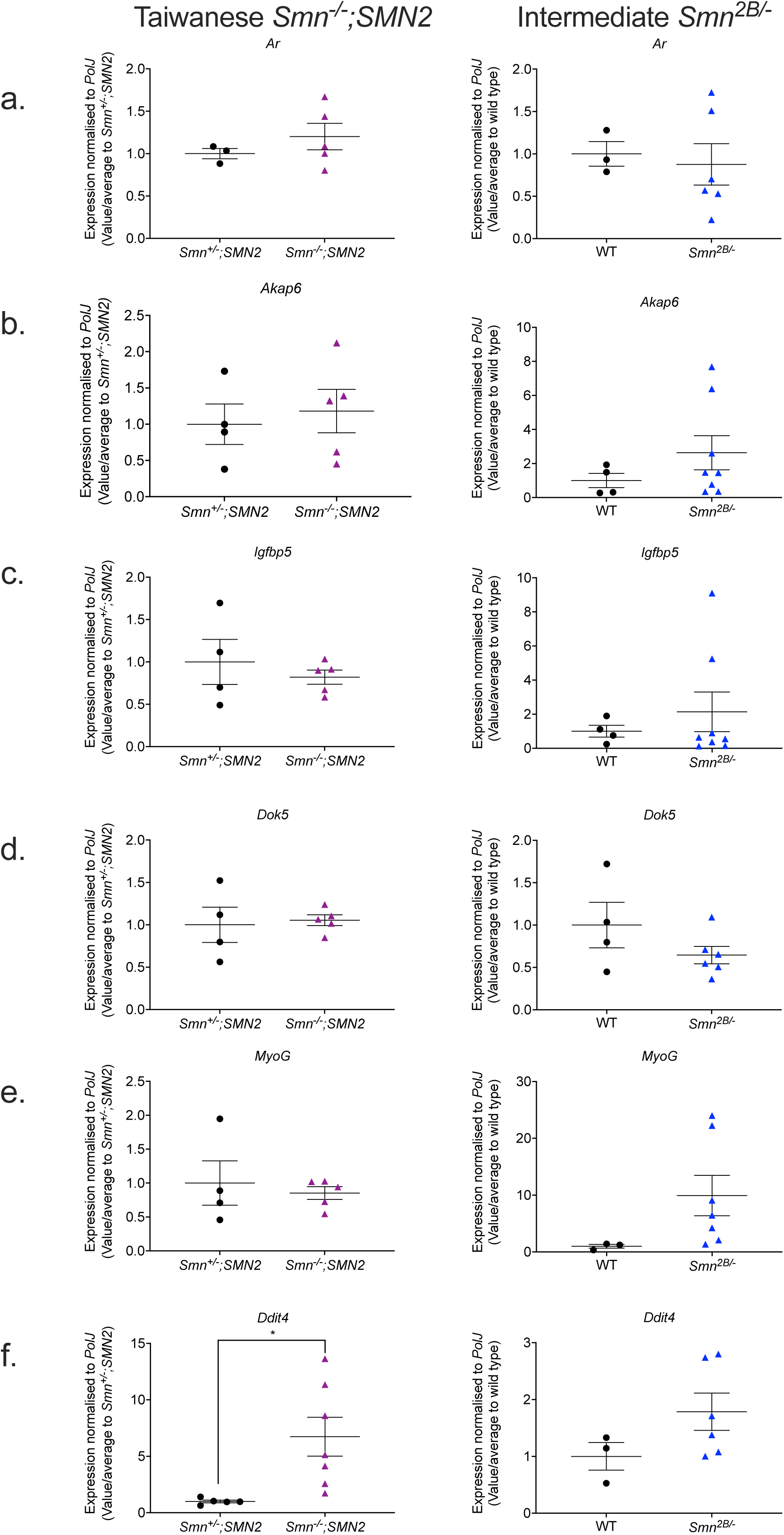
The oxandrolone target gene, *Ddit4*, is significantly upregulated in the skeletal muscle of severe *Smn^-/-^;SMN2* SMA mice. qPCR analysis of mRNA levels for predicted oxandrolone target genes **a.** *Ar*, **b.** *Akap6*, **c.** *Igfbp5*, **d.** *Dok5,* **e.** *MyoG* and **f.** *Ddit4* in the harvested triceps of untreated P7 Taiwanese *Smn^-/-^;SMN2* SMA mice (violet) and healthy *Smn^+/-^;SMN2* controls (black) (left panel) and symptomatic untreated P19 intermediate *Smn^2B/-^* SMA mice (blue) and wild type (C57BL/6J background) controls (black) (right panel). Data are shown as scatter plot that represents mean ± SEM error bars; n = 4-7 animals per experimental group, unpaired t-test, \**p* <0.05. *Smn^-/-^;SMN2 Ar*: *p* = 0.38; *Smn^-/-^;SMN2 Akap6*: *p* = 0.68; *Smn^-/-^;SMN2 Igfbp5*: *p* = 0.49; *Smn^-/-^;SMN2 Dok5*: *p* = 0.79; *Smn^-/-^ ;SMN2 MyoG*: *p* = 0.64; *Smn^-/-^;SMN2 Ddit4*: *p* = 0.02; *Smn^2B/-^ Ar*: *p* = 0. 75; *Smn^2B/-^ Akap6*: *p* = 0. 29; *Smn^2B/-^ Igfbp5*: *p* = 0.52; *Smn^2B/-^ Dok5*: *p* = 0.19; *Smn^2B/-^ MyoG*: *p* = 0.15, *Smn^2B/-^ Ddit4*: *p* = 0.16.

### *In vitro* oxandrolone treatment prevents canonical atrophy in C2C12 myotubes independently of the predicted Smn-independent targets

Similar to our metformin *in vitro* studies, we wanted to evaluate whether reduced SMN levels or atrophy influenced the expression of the predicted oxandrolone target genes in SMA skeletal muscle. Although none of the target genes were affected in the Smn-depleted C2C12 myoblasts (Figure 8.a), we found that Smn KD triggered a significant upregulation of *Dok5* only in C2C12 myotubes (Figure 8.b), suggesting that the expression of this gene may be Smn-dependent. Nevertheless, the expression of the majority of the predicted oxandrolone target genes was Smn- independent.

**Figure 8.**
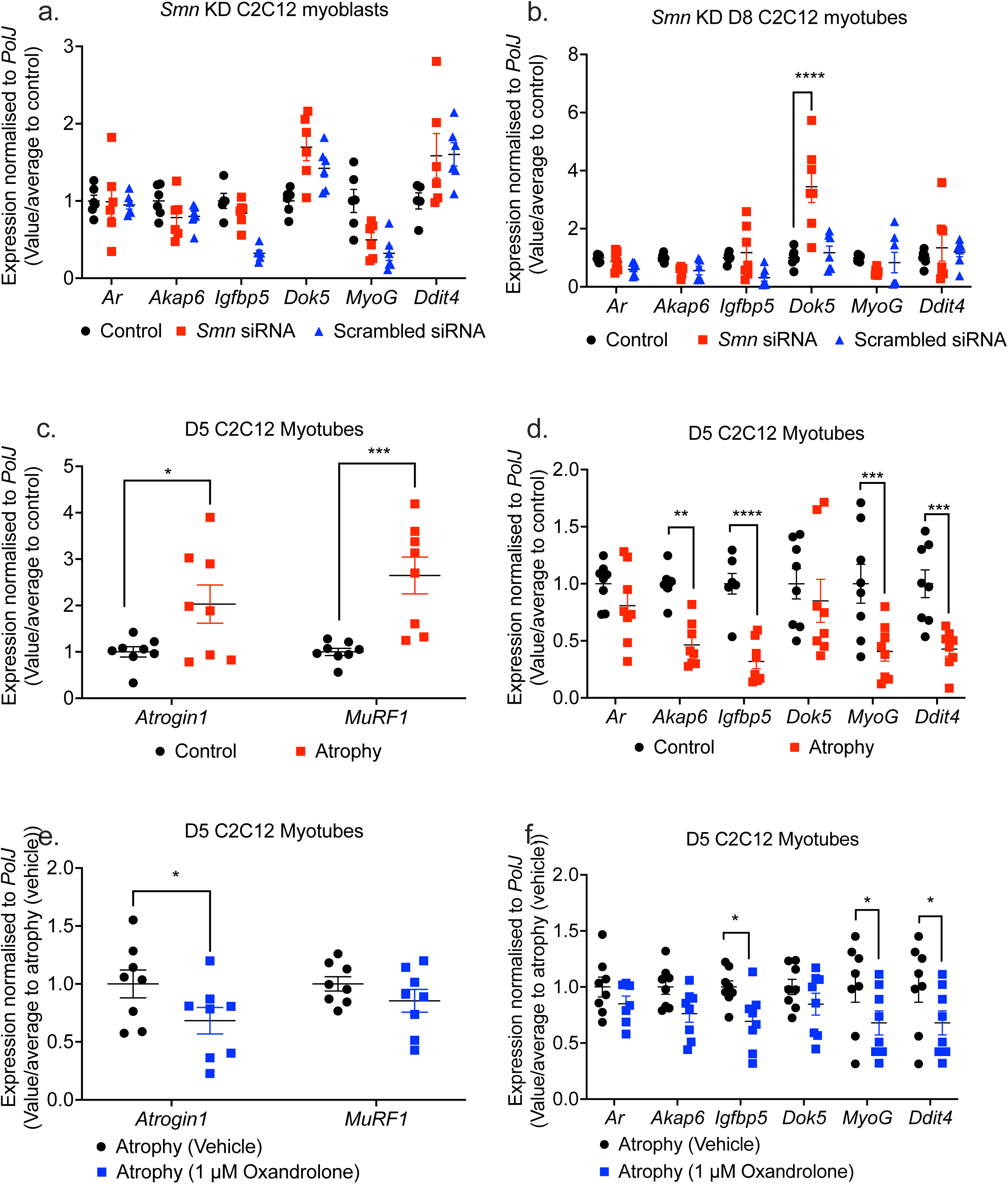
Oxandrolone target genes are pre-dominantly SMN-independent in SMA muscle C2C12 cellular model. *Smn* siRNA knockdown (red) was performed for **a.** 48 hours in C2C12 myoblasts and **b.** every 48 hours throughout differentiation in D8 C2C12 myotubes. mRNA expression of oxandrolone target genes *Ar*, *Akap6*, *Igfbp5*, *Dok5, MyoG* and *Ddit4* was measured by qPCR and compared to non- transfected (black) and scrambled siRNA transfected (blue) controls. D5 C2C12 myotubes were serum-starved for 24 hours to induce canonical atrophy (red). mRNA expression of **c.** atrogenes *Atrogin-1* and *MuRF-1* and **d.** oxandrolone target genes *Ar*, *Akap6*, *Igfbp5*, *Dok5, MyoG* and *Ddit4* was measured by qPCR and compared to non-starved myotubes (black). Serum-starved D5 C2C12 myotubes were treated with1 μM oxandrolone for 24 hours (blue) to evaluate mRNA expression via qPCR of **e.** atrogenes *Atrogin-1* and *MuRF-1* and **f.** oxandrolone target genes *Ar*, *Akap6*, *Igfbp5*, *Dok5, MyoG* and *Ddit4* compared to serum-starved absolute ethanol vehicle treated control (black). Data are shown as scatter graphs that represent mean ± SEM error bars; n = 4 samples per group across two independent experiments. Two-way ANOVA followed by uncorrected Fisher’s least significant difference (LSD). **a.** F = 5.45; **b.** F = 6.87; **c.** F = 1.1; **d.** F = 2.03; **e.** F = 0.72; **f.** F = 0.36, \**p* < 0.05, \*\**p* < 0.01, \*\*\**p* < 0.001, \*\*\*\**p* < 0.0001.

We next wanted to evaluate the ability of oxandrolone to attenuate canonical atrophy in serum-deprived C2C12 myotubes ^87^. However, in this case we performed the treatments in D5 C2C12 myotubes instead of D8, as although oxandrolone was non-toxic (Figures S8), it elicited a greater androgen *Ar* response at the earlier differentiation stage (Figure S9). Following confirmation of muscle atrophy in D5 C2C12 myotubes via significant *Atrogin-1* and *MuRF1* upregulation (Figure 8.c), we observed that the expression of the predicted oxandrolone target genes *Akap6*, *Igfbp5*, *MyoG* and *Ddit4* was significantly downregulated in serum-deprived D5 C2C12 myotubes (Figure 8.d).

Interestingly, we found that 24-hour treatment with 1 μM oxandrolone attenuated canonical muscle atrophy in these serum-starved D5 C2C12 myotubes as shown by significant downregulation of *Atrogin-1* (Figure 8.e). However, we observed that *Igfbp5*, *MyoG* and *Ddit4* were further downregulated by the 1 μM oxandrolone treatment (Figure 8.f), suggesting that oxandrolone’s effects on atrophy are linked to effectors independent of the predicted target genes. Overall, our *in vitro* studies have shown that although the expression of the predicted oxandrolone target genes is Smn-independent, they are not involved in oxandrolone’s ameliorative effects on canonical atrophy in C2C12 myotubes.

### Oxandrolone treatment improves survival in *Smn^2B/-^* SMA mice

We next assessed the impact of oxandrolone in SMA mice. We initially tested preliminary treatment regimens of 1 – 8 mg/kg/day starting from P5 or P8 in *Smn^2B/-^* SMA and *Smn^2B/+^* healthy mice (data not shown), based on previous studies in models of spinal cord injury (SCI) ^107^ and burn injury ^108^. We also stopped oxandrolone treatments at P21 as previous research has shown that shorter oxandrolone treatments are more effective ^107^. These pilot studies allowed us to identify the optimal dosing regimen of 4 mg/kg/day oxandrolone treatment from P8 to P21, which significantly improved the median survival of *Smn^2B/-^* SMA mice (Figure 9.a).

**Figure 9.**
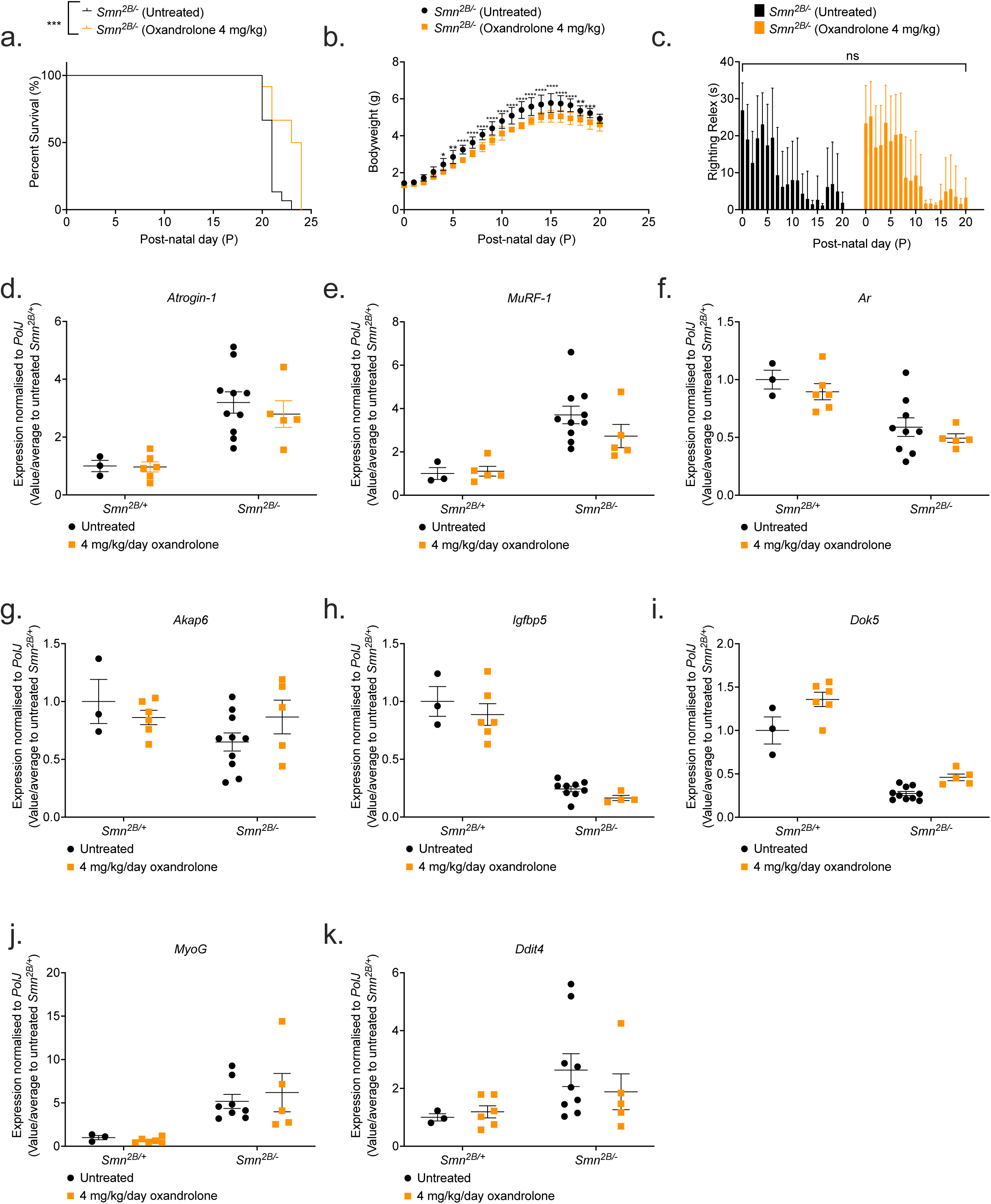
4 mg/kg/day oxandrolone treatment partially improves survival in *Smn^2B/-^* SMA mice. All treated animals received a daily dose of oxandrolone (4 mg/kg/day, suspended in 0.5% CMC) by gavage starting at P8. **a.** Survival curves of untreated (n = 15, 21 days median survival, black) and 4 mg/kg/day oxandrolone-treated (n = 12, 24 days median survival, orange) *Smn^2B/-^* SMA mice. Kaplan-Meier survival curve shown with log rank (Mantel-Cox) test, \*\*\**p* = 0.0006. **b.** Daily weights of untreated (n = 15, black) and 4 mg/kg/day oxandrolone-treated (n = 12, orange) *Smn^2B/-^* SMA mice. Data represented as mean ± SEM error bars; Two-way ANOVA followed by a Sidak’s multiple comparison test, F = 610.8, df = 519, \**p* < 0.05, \*\**p* < 0.01, \*\*\**p* < 0.001, \*\*\*\**p* < 0.0001. **c.** Daily righting reflex test for motor function activity up to a 30 second maximum time point in untreated (n = 15, black) and 4 mg/kg/day oxandrolone-treated (n = 12, orange) *Smn^2B/-^* SMA mice. Data are shown as bar chart with mean ± SEM error bars; unpaired T-test, ns = not significant, *p* = 0.775. qPCR analysis of mRNA levels for atrogenes **d.** *Atrogin-1* and **e.** *MuRF1* and predicted target genes **f.** *Ar,* **g.** *Akap6,* **h.** *Igfbp5,* **i.** *Dok5,* **j.** *MyoG,* and **k.** *Ddit4* in the triceps muscle from untreated (black) and 4 mg/kg/day oxandrolone-treated (orange), P19 *Smn^2B/+^* healthy and *Smn^2B/-^* SMA mice. Data are shown as bar chart with scatter graph represented as mean ± SEM error bars, n = 4 animals per group; two way ANOVA with Tukey’s multiple comparisons test, *Atrogin-1* (F = 0.1914), *MuRF1* (F = 1.214), *Ar* (F = 0.003), *Akap6* (F = 2.40), *Igfbp5* (F = 0.06), *Dok5* (F = 1.72), *MyoG* (F = 0.29) and *Ddit4* (F = 0.64), \**p* < 0.05, \*\**p* < 0.01.

However, we found that the body weight of 4 mg/kg/day oxandrolone-treated *Smn^2B/-^* SMA mice was significantly lower compared to their untreated counterparts (Figure 9.b), which is most likely due to the intrinsic smaller sizes of the randomly assigned litters, as demonstrated by the difference in weight starting 4 days prior to initial treatment (Figure 9.b). In terms of motor function, we observed no significant difference in righting reflex between untreated and oxandrolone-treated SMA animals (Figure 9.c). Furthermore, we identified no impact of vehicle treatment on survival, weight, and righting reflex in *Smn^2B/-^* SMA mice (Figures S10).

In the *Smn^2B/+^*healthy mice, although 4 mg/kg/day oxandrolone had no effect on survival or motor function in treated animals (Figures S11.a-b), we did observe a significant decrease in bodyweight starting from P9, one day after initial treatment (Figure S11.c), suggesting that oxandrolone may have impacted growth.

Nevertheless, our results demonstrated that although 4 mg/kg/day oxandrolone treatment improved survival in *Smn^2B/-^* SMA mice, its effect on survival was still minor compared to prednisolone ^27^, suggesting that oxandrolone is not a suitable substitute as an SMA skeletal muscle therapy.

### Oxandrolone did not impact the expression of the predicted target genes or muscle pathology markers

The improved 3-day survival in 4 mg/kg/day oxandrolone-treated *Smn^2B/-^* SMA mice led us to evaluate whether this beneficial impact was related to targeting muscle pathologies. Thus, we evaluated oxandrolone’s effects on the expression of dysregulated molecular markers associated with the SMA hallmark pathology of muscle atrophy (*Atrogin-1* and *MuRF-1*) in the triceps from P19 late symptomatic, untreated and 4 mg/kg/day oxandrolone-treated *Smn^2B/-^* SMA and *Smn^2B/+^* healthy mice, 2 hours after final treatment. We observed no significant reduction in elevated *Atrogin-1* or *MuRF-1* gene expression levels by oxandrolone in the *Smn^2B/-^* SMA cohort (Figures 9.d-e), suggesting that oxandrolone did not attenuate muscle atrophy.

We next evaluated the effect of oxandrolone on the expression of the predicted target genes in the same P19 untreated and 4 mg/kg/day oxandrolone-treated *Smn^2B/-^* SMA and *Smn^2B/+^* healthy mice. We found that oxandrolone did not significantly impact the predicated target genes in the triceps from the *Smn^2B/-^* SMA mice (Figures 9.f-k). However, we did observe that 4 mg/kg/day oxandrolone treatment significantly upregulated *Dok5* expression in the *Smn^2B/+^* healthy mice (Figure 9.i). Nevertheless, the pattern observed suggests that oxandrolone did not impact any of the predicted genes in the muscle from *Smn^2B/-^* SMA mice.

Overall, our data shows that oxandrolone did not have an efficient effect on the predicted target genes. Furthermore, its inability to ameliorate muscle atrophy marker dysregulation in SMA skeletal muscle, suggests that improved survival in the *Smn^2B/-^* SMA mice by oxandrolone may be independent of targeting skeletal muscle pathologies.

### Both metformin and oxandrolone drug candidates attenuate neuromuscular phenotypes in the *C. elegans* severe SMA model

We next wanted to investigate our drug candidates in a separate SMA model to assess whether they could attenuate neuromuscular dysfunctions in a distinct species. For this purpose, we used the *C. elegans smn-1(ok355)* invertebrate model ^63^, based on shared conservation of the SMN protein with vertebrate species ^109^ and the well described phenotypic defects of larval lethality (reduced survival) and impaired neuromuscular function in pharyngeal pumping for feeding and locomotion ^63, 65^. For metformin, administration at higher doses of 50 mM partially ameliorated multiple phenotypes only in *C. elegans smn-1(ok355)* including pharyngeal pumping defects and the locomotory defect of number of reversals (Figures 10.a-b), however only the lower dose of 1 mM metformin significantly ameliorated paralysis times (Figures 10.c). On the other hand, oxandrolone across 1-50 μM doses significantly ameliorated pharyngeal pumping defects only (Figure 10.d), with no significant effect on locomotory defects of reversal and paralysis times (Figures 10.e-f), suggesting an improvement in muscular activity in the pharynx region. Overall, given SMN conservation across species, oxandrolone could improve neuromuscular defects across vertebrate and invertebrate models of SMA.

**Figure 10.**
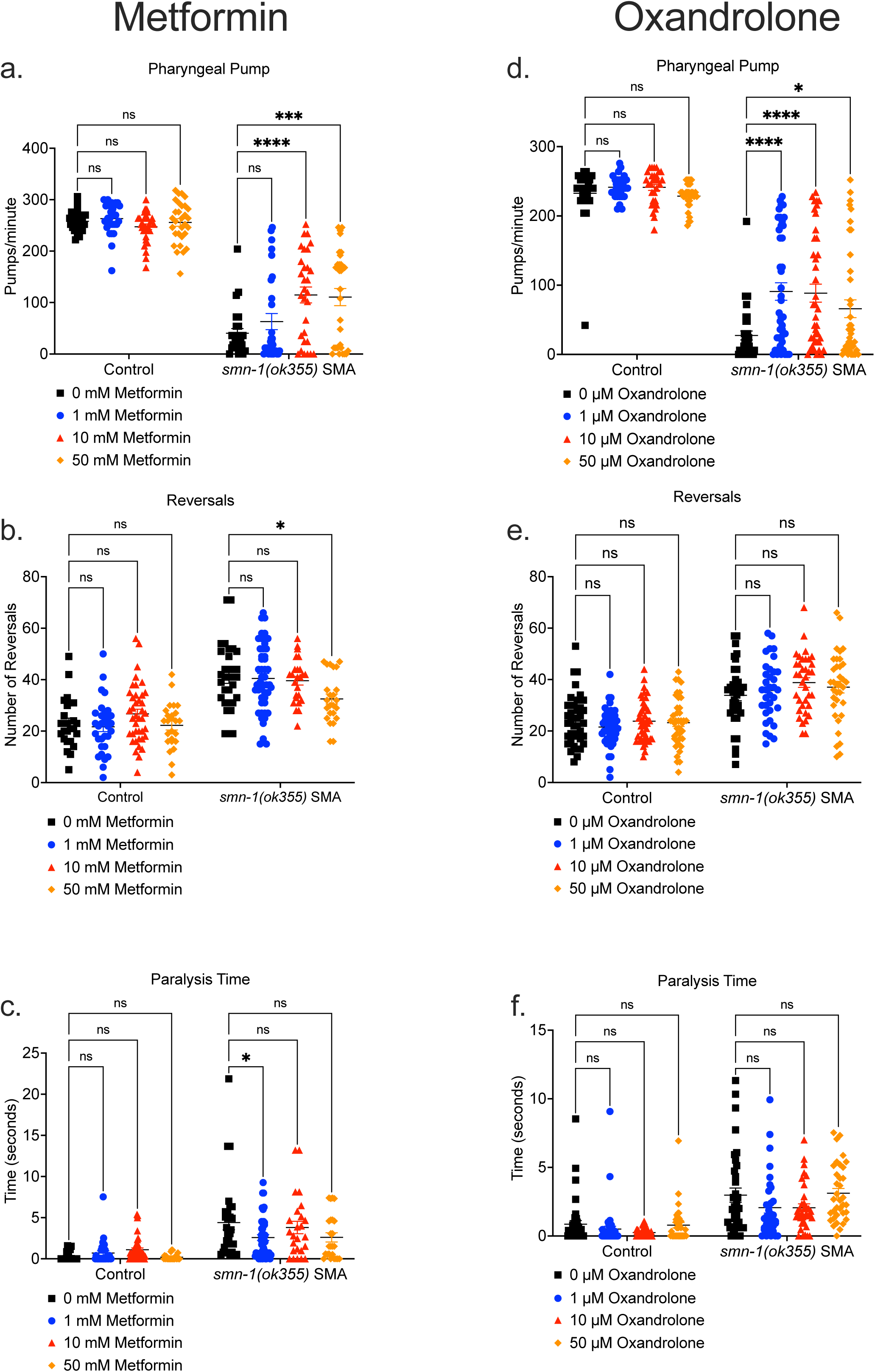
Metformin and oxandrolone partially ameliorates neuromuscular defects in the severe SMA *C. elegans* model. Day 3 *C. elegans smn-1(ok355)* SMA homozygotes and *smn-1(ok355)I/hT2* control heterozygotes were maintained at 20 °C on nematode growth medium (NGM) plates seeded with Escherichia coli OP50 bacteria. In metformin conditions, the NGM contained metformin concentrations of 0 (black square), 1 (blue circle), 10 (red triangle) and 50 mM (orange diamond) respectively. **a.** Pharyngeal pumping rates (pumps/minute) defined as grinder movement in any axis at 175 frames/10 seconds. Locomotion assays were filmed at 15 frames/sec and quantified for 5 minutes for **b.** reversals and **c.** paralysis times in the *C. elegans* groups. In oxandrolone conditions, the NGM contained oxandrolone concentrations of 0 (black square), 1 (blue circle), 10 (red triangle) and 50 μM (orange diamond) respectively. **d.** Pharyngeal pumping rates (pumps/minute) defined as grinder movement in any axis at 175 frames/10 seconds. Locomotion assays were filmed at 15 frames/sec and were quantified for 5 minutes for **e.** reversals and **f.** paralysis times in the *C. elegans* groups. Data are shown as scatter plot that represents mean ± SEM error bars; n > 25 animals per experimental group across three independent trials; Two-way ANOVA with Tukey’s multiple comparisons test, **a.** F = 459.2, **b.** F = 108.6, **c.** F = 52.59, **d.** F = 501.8, **e.** F = 121.1, **f.** F = 57.06, ns = not significant \**p* < 0.05, \*\**p* < 0.01, \*\*\**p* < 0.001.

## DISCUSSION

The goal of this study was to use a transcriptomics-based drug repositioning strategy to identify clinically approved drug candidates that could emulate prednisolone’s beneficial effects in SMA skeletal muscle and life expectancy ^27^, without the risks associated with long-term GC exposure^39^.

Our major finding was the observation that prednisolone treatment restored specific gene sets associated with key pathological SMA pathways such as FoxO signalling ^66^, p53 signalling ^67^, AMPK signalling ^68^, mitophagy ^69^, circadian rhythm ^70^, PPAR signalling ^71^, and autophagy ^72^ in *Smn^-/-^;SMN2* SMA skeletal muscle. Although these pathway results highlight prednisolone’s efficacy in improving skeletal muscle health, it should be noted that our transcriptomic data cannot distinguish whether these restorations are targeted directly by prednisolone or a consequence of improved muscle health. Nevertheless, our transcriptomic data complemented the prior phenotypic data of prednisolone’s potential as a second-generation SMA therapy ^27^.

Despite a multitude of promising compounds identified that could be investigated in future studies, one of the findings of our study was that neither of our chosen orally bioavailable drug candidates, metformin and oxandrolone, reproduced prednisolone’s previously reported effect on muscle growth and survival in SMA mice ^27^. For metformin, we observed that both 200- and 400- mg/kg/day doses counterintuitively exacerbated *Prkag3* downregulation in *Smn^2B/-^* SMA muscle instead of the predicted upregulation, which could have negative consequences since *Prkag3^-/-^* null mice presented metabolic and mitochondrial dysregulation ^110, 111^ similar to those reported in SMA patients ^69, 93^ and models ^89, 90^. More surprising was the reduced survival of *Smn^2B/-^* SMA mice treated with 400 mg/kg/day metformin, which we potentially linked to possible hypoglycaemic shock and/or dysregulation of neuronal mitochondrial components. On the other hand, although various metformin doses ameliorated neuromuscular dysfunction in our SMA *C. elegans* model, the negative effects observed in *Smn^2B/-^* SMA mice could be to due vertebrate vs invertebrate systems. Curiously, previous research into AMPK agonism via 500 mg/kg AICAR treatment in the *Smn/17* SMA mice contrasts our metformin data, as they showed improved skeletal muscle health and no exacerbations in neuronal dysfunction ^68^. One explanation for metformin and AICAR’s conflicting results in SMA mice could be related to differential pharmacological activities. Indeed, AICAR is an AMP analogy with a low BBB penetrability ^112^ that directly binds to the γ-AMPK isoform ^113, 114^, whilst metformin can rapidly penetrate the BBB ^98^ and has been associated with direct and indirect AMPK activation ^97^, an example of the latter involving mitochondrial respiratory complex 1 inhibition ^96^. With emerging evidence of naturally low mitochondrial respiratory complex 1 activity in SMA motor neurons ^115^ and SMA pathway profiles being tissue-dependent ^72^, one theory could be that the 400 mg/kg/day metformin dose exacerbated mitochondrial respiratory complex 1 inhibition in SMA motor neurons. However, future studies would be needed to verify this proposed model. Nonetheless, our findings could be important for clinical drug safety, as with reported co-morbidities of diabetes in certain SMA patients ^116, 117^, our pre-clinical data suggests lower metformin doses or non-biguanide drugs may be important to manage diabetes and not risk primary pathologies in SMA patients.

For oxandrolone, our mouse data showed that 4 mg/kg/day treatment from P8 partially improved survival in *Smn^2B/-^* SMA mice, although not to the same extent observed with prednisolone ^27^. In addition, we identified that the lower dose of 1 μM oxandrolone *in vitro* attenuated canonical atrophy in C2C12 myotubes, whilst *in vivo* oxandrolone attenuated neuromuscular dysfunction in severe SMA *C. elegans* model, suggesting that in both our SMA vertebrate and invertebrate species oxandrolone may be a beneficial SMN-independent treatment option. However, one factor that we did not account for in our oxandrolone investigations was gender specificity. Although the clinical literature has shown oxandrolone has minimal androgenous effects, studies with *Ar* KO rodent models have demonstrated that Ar absence does not have the same impact on female muscle size compared to males ^118^. Furthermore, it is suggested that intramuscular Ar content may have a stronger influence on hypertrophy than peripheral androgen levels ^119^. Thus, we cannot conclude whether oxandrolone’s treatment efficacy compared to prednisolone was due to gender-specific differences.

Despite our study’s limitations, it highlighted refinements for future *in silico* SMA drug repositioning studies. Compared to a previous study that successfully identified and validated harmine’s therapeutic potential in SMA muscle ^42^, ours did not include proteomics. The absence of proteomics can be a caveat for drug studies as transcript levels alone do not proportionally reflect drug-protein interactions, abundance and translational modifications ^120, 121^. However, a limitation of both transcriptomics and proteomics approaches is that they cannot bridge drug-pathway interactions with disease phenotypes, as demonstrated by a recent proteomics analysis of Spinraza® treated type 2 and 3 SMA patients that could not correlate protein profiles with functional improvements ^122^. Thus, implementation of metabolomics may be beneficial for linking pathway perturbations with metabolites associated with disease and stages of muscle atrophy ^123^. To the best of our knowledge, this three-pronged multi-omics approach has not previously been used in SMA drug repositioning, however it has been successful in the identification of 200 biomarker candidates for SMA ^124^.

Another consideration is systemic effects of the drugs as seen with the enhanced lethality of the 400 mg/kg/day metformin’s dose in *Smn^2B/-^*SMA mice being linked to hypoglycaemia and mitochondrial dysfunction in neuronal tissue. Indeed, adverse systemic risks were also found with a previous multi-omics drug repositioning study for SMA muscle that identified the development of tremors in harmine-treated *Smn^2B/-^* SMA mice ^42^. With tissue-dependent differences in conserved pathways in SMA ^72^, future omics studies could integrate data from both neuronal and skeletal muscle to minimize systemic adverse risks.

Even with these refinements, future drug repositioning studies for SMA skeletal muscle may need to consider replacing bulk RNA-Seq. Skeletal muscle fibers are comprised of a myriad of different muscle fiber and cell types, alongside non-muscular interconnecting tissues such as neurons, tendons, adipose, immune cells and capillaries ^125, 126^, which altogether does not truly represent the transcriptomic profiles for distinct skeletal muscle cells. Indeed, our significant KEGG pathway results included those associated exclusively with neuronal, immune, and capillary cells such as glioma, atherosclerosis and Th17 cell differentiation. With alterations in fiber type composition ^91, 127^ and satellite cell dysregulation evidenced in SMA muscle ^23^, an emerging alternative to predict drug candidates that target dysregulated SMA pathways in these muscle types would be single-cell (scRNA-Seq) and/or single nuclear RNA-Seq (snRNA-Seq) ^126^, which have already been useful tools in other muscular disorders such as DMD ^128, 129^. In addition, snRNA-Seq could have further benefits to narrow in on muscle regions such as nuclei located near the NMJ, since myonuclei display spatial and temporal expression pattern heterogeneity in multi-nucleated myofibers ^126^.

Although the drug candidate’s metformin and oxandrolone did not emulate prednisolone’s beneficial effects in SMA to the extent previously reported, our transcriptomics-drug repositioning approach did better define prednisolone’s activity in SMA muscle and provided a list of potential candidates for future pre-clinical SMA drug repositioning studies. Furthermore, our study highlights important refinements for future SMA drug repositioning studies.

## Data Availability Statement

The datasets presented in this study can be found in the following online repository: NCBI BioProject, accession ID: PRJNA972323.

## AUTHOR CONTRIBUTIONS

Conceptualization: J.M.H, M.B; Methodology: J.M.H, P.P.T, M.D, L.M.W, P.C, D.P.T & M.B; Formal Analysis: J.M.H, P.P.T, M.D, D.P.T & M.B; Investigation: J.M.H, E.M, E.R.S, O.C, P.P.T, M.D, M.O, L.M.W, P.C, & M.B; Software: J.M.H & D.P.T; Visualization: J.M.H; Resources: M.D, P.C, S.D, M.J.A.W, D.P.T & M.B; Writing – Original Draft: J.M.H & M.B; Writing – Review & Editing: J.M.H, E.M, E.R.S, O.C, P.P.T, M.D, M.O, L.M.W, P.C, S.D, M.J.A.W, D.P.T, & M.B; Supervision: D.P.T & M.B; Project Administration: M.B; Funding Acquisition: M.B

## FUNDING

J.M.H. was funded by a Ph.D. studentship from the Keele University of School of Medicine. E.M. is supported by an Academy of Medical Sciences grant (SBF006/1162). E.R.S. was funded by a MDUK Ph.D. studentship (18GRO-PS48-0114). O.C. is supported by a Ph.D. studentship from the Republic of Turkey Ministry of National Education. M.B. was funded by SMA Angels Charity. P.C. is supported by the Deutsche Muskelstiftung and the European Union’s Horizon 2020 research and innovation programme under the Marie Sklodowska-Curie grant agreement No 956185.

## ACKNOWLEDGEMENTS

We would like to thank the personnel at the Biomedical Sciences Unit at both Keele University and University of Oxford. Many thanks go to members of the Combined Inherited Neuromuscular Disorders meetings for helpful discussions. For the RNA-Sequencing we are grateful for the aid and resources of Research Core Unit Transcriptomics at Hannover Medical School. *C. elegans* strains were provided by the *Caenorhabditis elegans* consortium (CGC), which is funded by NIH Office of Research Infrastructure Programs (P40 OD010440). Research in the M.D. lab has been funded by the Wellcome Trust and Royal Society. A special thanks goes out to Advaita for their technical support of iPathwayGuide and to CureSMA, SMA Europe and the Biochemical Society, UK for providing podium presentation opportunities for preliminary data of this project.

## SUPPLEMENTARY FIGURE LEGENDS

**Figure S1. “Condition” and “Treatment” groups share distinct transcriptomic patterns for a specific sub-set of genes in prednisolone-treated *Smn^-/-^;SMN2* SMA mice and untreated *Smn^-/-^;SMN2* SMA and *Smn^+/-^;SMN2* healthy mice.**

*Smn^-/-^;SMN2* SMA and *Smn^+/-^;SMN2* healthy mice received prednisolone treatment (5 mg/kg gavage every 2 days) from P0. The triceps was harvested from P7 untreated and prednisolone-treated *Smn^-/-^;SMN2* SMA and *Smn^+/-^;SMN2* healthy mice for RNA isolation and library preparation for RNA-Sequencing. Differential gene expression analysis was performed by DESeq2 v2.11.40.2 with study design set to “condition and “treatment”. Principal component analysis based on transcriptomic profiles between **a.** P7 untreated *Smn^+/-^;SMN2* (red, n=3) and untreated *Smn^-/-^;SMN2* (blue, n=3) mice. **b.** P7 untreated *Smn^-/-^;SMN2* (green, n=3) and prednisolone-treated *Smn^-/-^;SMN2* (blue, n=2) SMA mice.

**Figure S2. Bioinformatic identification of shared target genes between metformin and prednisolone in SMA skeletal muscle.**

**a.** The pathway diagram contains proteins within the FOXO signalling pathway (KEGG: 04068) encoded by the predicted differentially expressed genes from prednisolone vs untreated *Smn^-/-^;SMN2* SMA skeletal muscle. The AMPK protein (yellow circle) represents the AMPK-γ3 isoform gene *Prkag3*. Its activity on the iPathwayGuide interactive server highlights that it can be targeted by metformin via a built-in KEGG Drugs database. The red lines represent coherent cascades, which supports the consistency of the transcriptomic data and published pathway activity. In the FOXO signalling pathway, activation of AMPK represented by *Prkag3* coherently downregulates the FOXO protein (green circle), which represents *Foxo1*, *Foxo3*, *Foxo4*, and *Foxo6* isoforms. The highest logFC patterns are shown in dark red and lowest in dark blue as indicated in the legend value box. Graph generated in iPathwayGuide (Advaita). **b.** Differential gene expression pattern by logFC (Y-axis) of predicted metformin target genes *Prkag3*, *Foxo1*, *Foxo3*, *Foxo4* and *Foxo6* based on transcriptomic data from prednisolone vs untreated *Smn^-/-^;SMN2* SMA skeletal muscle. Upregulated genes above X-axis are highlighted in red and downregulated genes below X-axis are highlighted in blue. The box and whisker plot on Y-axis represents 1^st^ quartile, median and 3^rd^ quartile. Graph generated in iPathwayGuide (Advaita). **c.** Heatmap visualization for predicted metformin targets *Prkag3*, *Foxo1*, *Foxo3*, *Foxo4* and *Foxo6* (Log2FC > 0.6; FDR < 0.05) between untreated *Smn^+/-^;SMN2* healthy mice (left), untreated *Smn^-/-^;SMN2* SMA mice (centre) and prednisolone-treated *Smn^-/-^;SMN2* SMA mice (right). Colour key represents log2FC for upregulated (red) and downregulated (blue) genes. Heatmap was generated by Heatmap v2.1.1+galaxy1.

**Figure S3. Smn knockdown in C2C12 myoblasts and myotubes via *Smn* siRNA transfection.** *Smn* siRNA knockdown (red) was performed for **a.** 48 hours in C2C12 myoblasts and **b.** every 48 hours throughout differentiation in D8 C2C12 myotubes. *Smn* knockdown of mRNA levels was confirmed by qPCR and compared to non-transfected (black) and scrambled siRNA control (blue) groups. Data are shown as scatter graph that represent mean ± SEM error bars; n = 4 samples per group across two independent experiments. One-way ANOVA followed by Tukey’s multiple comparisons test. C2C12 myoblasts F = 22.01; D8 C2C12 myotubes F = 115.4. \*\*\*\**p* <0.0001.

**Figure S4. Both physiological and supraphysiological metformin concentrations affect the expression of specific predicted target genes in C2C12 myoblasts and myotubes. a.** C2C12 myoblasts and **b.** D8 C2C12 myotubes were treated with a range of physiological (30 μM (red) and 60 μM (green)) and supraphysiological (1 mM (brown) and 2 mM (blue)) metformin concentrations for 24 hours against a PBS vehicle control (black) to evaluate the mRNA expression via qPCR of predicted target genes *Prkag3*, *Foxo1*, *Foxo3*, *Foxo4* and *Foxo6*. Data are shown as bar charts with scatter graph that represent mean ± SEM error bars; n = 4 samples per group across two independent experiments. Two-way ANOVA followed by uncorrected Fisher’s least significant difference (LSD). C2C12 myoblasts F = 7.822; D8 C2C12 myotubes F = 12.17. \**p* <0.05, \*\**p* <0.01, \*\*\**p* <0.001, \*\*\*\**p* <0.0001.

**Figure S5. 0.9% saline vehicle treatment had no effect on survival or phenotype in *Smn^2B/-^* SMA and *Smn^2B/+^* healthy mice.**

All treated animals received a daily dose of 0.9% physiological saline vehicle by gavage starting at P5**. a.** Survival curves of untreated (n = 13, black) and 0.9% saline vehicle-treated (n = 14, green) *Smn^2B/-^*SMA mice. Kaplan-Meier survival curve shown with log rank (Mantel-Cox) test, ns = not significant, *p* = 0.0653. **b.** Daily weights of untreated (n = 13, black) and 0.9% saline vehicle-treated (n = 14, green) *Smn^2B/-^*SMA mice. Data represented as mean ± SEM error bars; Two-way ANOVA followed by a Sidak’s multiple comparison test, F = 454.6, df = 501 **c.** Daily righting reflex test for motor function activity up to a 30 second maximum time point in untreated (n = 13, black) and 0.9% saline vehicle-treated (n = 14, green) *Smn^2B/-^* SMA mice. Data are shown as bar chart with mean ± SEM error bars; unpaired T-test, ns = not significant, *p* = 0.6626**. d.** Survival curves of untreated (n = 16, black) and 0.9% saline vehicle-treated (n = 22, green) *Smn^2B/+^* healthy mice. Kaplan-Meier survival curve shown with log rank (Mantel-Cox) test, ns = not significant, *p* > 0.9999. **e.** Daily weights of untreated (n = 16, black) and 0.9% saline vehicle-treated (n = 22, green) *Smn^2B/+^* healthy mice. Data represented as mean ± SEM error bars; Two- way ANOVA followed by a Sidak’s multiple comparison test, F =328, df = 757. **f.** Daily righting reflex test for motor function activity up to a 30 second maximum time point in untreated (n = 16, black) and 0.9% saline vehicle-treated (n = 22, green) *Smn^2B/+^* healthy mice. Data are shown as bar chart with mean ± SEM error bars; unpaired T-test, ns = not significant, *p* = 0.9555.

**Figure S6. 200 and 400 mg/kg/day metformin had no negative effect on survival or phenotype in healthy *Smn^2B/+^*mice.**

All treated animals received a daily dose of metformin (either 200 or 400 mg/kg/day, diluted in 0.9% saline) by gavage starting at P5. **a.** Survival curves of untreated (n = 16, black) and 200 mg/kg/day metformin-treated (n = 20, red) *Smn^2B/+^* healthy mice. Kaplan-Meier survival curve shown with log rank (Mantel-Cox) test, ns = not significant, *p* > 0.9999. **b.** Daily weights of untreated (n = 16, black) and 200 mg/kg/day metformin-treated (n = 20, red) *Smn^2B/+^* healthy mice. Data represented as mean ± SEM error bars; Two-way ANOVA followed by a Sidak’s multiple comparison test, F = 549.5, df = 680. **c.** Daily righting reflex test for motor function activity up to a 30 second maximum time point in untreated (n = 16, black) and 200 mg/kg/day metformin- treated (n = 20, red) *Smn^2B/+^* healthy mice. Data are shown as bar chart with mean ± SEM error bars; unpaired T-test, ns = not significant, *p* = 0.9183**. d.** Survival curves of untreated (n = 16, black) and 400 mg/kg/day metformin-treated (n = 15, blue) *Smn^2B/+^*healthy mice. Kaplan-Meier survival curve shown with log rank (Mantel-Cox) test, ns = not significant, *p* > 0.9999. **e.** Daily weights of untreated (n = 16, black) and 400 mg/kg/day metformin-treated (n = 15, blue) *Smn^2B/+^*healthy mice. Data represented as mean ± SEM error bars; Two-way ANOVA followed by a Sidak’s multiple comparison test, F = 261.9, df = 435. **f.** Daily righting reflex test for motor function activity up to a 30 second maximum time point in untreated (n = 16, black) and 400 mg/kg/day metformin-treated (n = 15, blue) *Smn^2B/+^* healthy mice. Data are shown as bar chart with mean ± SEM error bars; unpaired T-test, ns = not significant, *p* = 0.9966.

**Figure S7. Oxandrolone is predicted to emulate the target patterns of prednisolone in the skeletal muscle (Triceps) of *Smn^-/-^;SMN2* SMA mice.**

**a.** Predicted model based on upstream regulator patterns predicted by iPathwayGuide in prednisolone vs untreated *Smn^-/-^;SMN2* SMA skeletal muscle. *Ar* upregulates downstream targets *Igfbp5* and *MyoG*, whilst it downregulates *Ddit4*. *Igfbp5* upregulates *Dok5* and *MyoG* upregulates *Akap6*. Upregulated genes shaded in red, downregulated genes shaded in blue. Downregulation of *Ddit4* based on previous published literature. **b.** Differential gene expression pattern by logFC (Y-axis) of predicted oxandrolone targets *Ddit4*, *Igfbp5*, *Ar*, *MyoG*, *Akap6* and *Dok5* based on transcriptomic data from prednisolone vs untreated *Smn^-/-^;SMN2* SMA skeletal muscle. Upregulated genes above X-axis are highlighted in red and downregulated genes below X-axis are highlighted in blue. The box and whisker plot on Y-axis represents 1^st^ quartile, median and 3^rd^ quartile. Graph generated in iPathwayGuide (Advaita). **c.** Heatmap visualization for predicted oxandrolone targets *Ar*, *MyoG*, *Igfbp5*, *Dok5*, *Akap6* and *Ddit4* (log2FC > 0.6; FDR < 0.05) between untreated *Smn^+/-^;SMN2* healthy mice (left), untreated *Smn^-/-^;SMN2* SMA mice (centre) and prednisolone-treated *Smn^-/-^;SMN2* SMA mice (right). Colour key represents log2FC for upregulated (red) and downregulated (blue) genes. Heatmap was generated by Heatmap v2.1.1+galaxy1.

**Figure S8. Low 1 μM oxandrolone treatment is non-toxic and does not impact proliferation in C2C12 myoblasts and myotubes.**

**a.** C2C12 myoblasts and **b.** D5 C2C12 myotubes were treated with 1 μM oxandrolone for 24 (purple) and 72 hours (orange) and compared to an absolute ethanol vehicle (24 hours red, 72 hours blue) in addition to untreated (black) and 1% Triton-X max lactate dehydrogenase (LDH) control (green). LDH levels in cell culture supernatant were measured by proportional fluorescence absorption readings (nm). C2C12 myoblasts were treated with 1 μM oxandrolone for **c.** 24 (orange) and **d.** 72 hours (purple) against an absolute ethanol vehicle (24 hours red, 72 hours blue) in addition to blank media (brown), blank cells (cyan), and untreated cells (black) controls. Absorption readings (nm) were measured from anti-BrDU antibody immunostained samples. Data are shown as bar charts that represent mean ± SEM error bars; n = 6 samples per group across one independent experiments; one-way ANOVA followed by Dunnett’s multiple comparisons test, **a.** F = 44.25, **b.** F = 3.092, **c.** F = 67.51, **d.** F = 64.71, \**p* < 0.05, \*\*\**p* < 0.001, \*\*\*\**p* < 0.0001, ns = not significant.

**Figure S9. D5 stage in C2C12 myotubes for oxandrolone to elicit an *Ar* gene response.**

**a.** C2C12 myoblasts were treated with a range of oxandrolone concentrations of 1 (blue), 10 (red) and 100 μM (green) for 24 hours and compared to an absolute ethanol (vehicle, black) to evaluate the mRNA expression via qPCR of predicted target genes *Ar*, *Dok5*, *Igfbp5*, *Akap6*, *MyoG* and *Ddit4*. Data are shown as a bar chart and scatter graph that represent mean ± SEM error bars; n = 4 samples per group across two independent experiments. Two-way ANOVA followed by uncorrected Fisher’s least significant difference (LSD), F = 1.693. \**p* <0.05, \*\**p* <0.01, \*\*\**p* <0.001, \*\*\*\**p* <0.0001. **b.** D8 C2C12 myotubes were treated with a range of oxandrolone concentrations of 1 (blue), 10 (red) and 100 μM (green) for 24 hours and compared to an absolute ethanol (vehicle, black) to evaluate the mRNA via qPCR of predicted target gene *Ar* and downstream targets *Igfbp5* and *MyoG*. Data are shown as bar chart and scatter graph that represent mean ± SEM error bars; n = 4 samples per group across two independent experiments. Two-way ANOVA followed by uncorrected Fisher’s least significant difference (LSD), F = 0.8708. **c.** D3 and **d.** D5 C2C12 myotubes were treated with 1 μM oxandrolone for 24 hours (blue) against an absolute ethanol (vehicle, black) to evaluate the mRNA levels via qPCR of *Ar*. Data are shown as scatter graphs that represent mean ± SEM error bars; n = 4 samples per group across one independent experiment; unpaired T-test D3 C2C12 myotubes *p* = 0.0203, D5 C2C12 myotubes *p* = 0.0480, \**p* <0.05. **e.** D5 C2C12 myotubes were treated with 1 μM oxandrolone (blue) for 24 hours against an absolute ethanol (vehicle, black) to evaluate the mRNA levels via qPCR of target genes *Dok5*, *Igfbp5*, *Akap6*, *MyoG* and *Ddit4*. Data are shown as bar charts with scatter graphs that represent mean ± SEM error bars; n = 4 samples per group across two independent experiments. Two-way ANOVA followed by uncorrected Fisher’s least significant difference (LSD), F = 0.7013.

**Figure S10. 0.5% carboxymethyl cellulose vehicle treatment had no effect on survival or phenotype in *Smn^2B/-^* SMA and *Smn^2B/+^* healthy mice.**

All treated animals received a daily dose of 0.5% carboxymethyl cellulose (CMC) vehicle by gavage starting at P8**. a.** Survival curves of untreated (n = 15, black) and 0.5% CMC vehicle-treated (n = 13, violet) *Smn^2B/-^*SMA mice. Kaplan-Meier survival curve shown with log rank (Mantel-Cox) test, ns = not significant, *p* = 0.4222. **b.** Daily weights of untreated (n = 15, black) and 0.5% CMC vehicle-treated (n = 13, violet) *Smn^2B-/-^* SMA mice. Data represented as mean ± SEM error bars; Two-way ANOVA followed by a Sidak’s multiple comparison test, F = 335 df = 511 **c.** Daily righting reflex test for motor function activity up to a 30 second maximum time point in untreated (n = 15, black) and 0.5% CMC vehicle-treated (n = 13, violet) *Smn^2B/-^* SMA mice. Data are shown as bar chart with mean ± SEM error bars; unpaired T-test, ns = not significant, *p* = 0.5602**. d.** Survival curves of untreated (n = 21, black) and 0.5% CMC vehicle-treated (n = 16, violet) *Smn^2B/+^* healthy mice. Kaplan-Meier survival curve shown with log rank (Mantel-Cox) test, ns = not significant, *p* > 0.9999. **e.** Daily weights of untreated (n = 21, black) and 0.5% CMC vehicle-treated (n = 16, violet) *Smn^2B/+^* healthy mice. Data represented as mean ± SEM error bars; Two-way ANOVA followed by a Sidak’s multiple comparison test, F = 377.3, df = 724. **f.** Daily righting reflex test for motor function activity up to a 30 second maximum time point untreated (n = 21, black) and 0.5% CMC vehicle-treated (n = 16, violet) *Smn^2B/+^* healthy mice. Data are shown as bar chart with mean ± SEM error bars; unpaired T-test, ns = not significant, *p* = 0.9638.

**Figure S11. 4 mg/kg/day oxandrolone decreased bodyweight in healthy *Smn^2B/+^* mice.**

All treated animals received a daily dose of oxandrolone (4 mg/kg/day, suspended in 0.5% CMC) by gavage starting at P8. **a.** Survival curves of untreated (n = 21, black) and 4 mg/kg/day oxandrolone-treated (n = 10, orange) *Smn^2B/+^* healthy mice. Kaplan-Meier survival curve shown with log rank (Mantel-Cox) test, ns = not significant, *p* > 0.9999. **b.** Daily weights of untreated (n = 21, black) and 4 mg/kg/day oxandrolone-treated (n = 10, orange) *Smn^2B/+^* healthy mice. Data represented as mean ± SEM error bars; Two-way ANOVA followed by a Sidak’s multiple comparison test, F = 395.5, df = 598, \**p* < 0.05, \*\**p* < 0.01, \*\*\**p* < 0.001, \*\*\*\**p* < 0.0001. **c.** Daily righting reflex test for motor function activity up to a 30 second maximum time point in untreated (n = 21, black) and 4 mg/kg/day oxandrolone-treated (n = 10, orange) *Smn^2B/+^* healthy mice. Data are shown as bar chart with mean ± SEM error bars; unpaired T-test, ns = not significant, *p* = 0.7865.

## SUPPLEMENTARY TABLES

**Table S1. RNA sequencing sample groups.**

**Table S2. Murine primers for qPCR.**

**Table S3. Significant differentially expressed genes (Log2FC > 0.6; FDR < 0.05) in skeletal muscle (triceps) between P7 untreated *Smn^-/-^;SMN2* SMA and *Smn^+/-^;SMN2* healthy mice.**

**Table S4. Significant differentially expressed genes (Log2FC > 0.6; FDR < 0.05) in skeletal muscle (triceps) between P7 prednisolone-treated vs untreated *Smn^-/-^;SMN2* SMA mice.**

**Table S5. Raw Counts for significant differentially expressed genes in skeletal muscle (triceps) between P7 prednisolone-treated vs untreated *Smn^-/-^;SMN2* SMA and untreated *Smn^+/-^;SMN2* healthy mice.**

**Table S6. Complete list of significant Gene Ontology (GO) Term (Biological Processes) for P7 prednisolone-treated vs untreated *Smn^-/-^;SMN2* SMA mice.**

**Table S7. Complete list of significant Gene Ontology (GO) Term (Molecular Functions) for P7 prednisolone-treated vs untreated *Smn^-/-^;SMN2* SMA mice.**

**Table S8. Complete list of significant Gene Ontology (GO) Term (Cell Components) for P7 prednisolone-treated vs untreated *Smn^-/-^;SMN2* SMA mice.**

**Table S9. List of drugs identified in iPathwayGuide from KEGGS drug database that target significantly impacted prednisolone pathways.**

**Table S10. List of predicted upstream regulators from P7 prednisolone-treated vs untreated *Smn^-/-^;SMN2* SMA mice used in DGIdb v.3. gene-drug target search.**

**Table S11. List of predicted agonist drugs from DGIdb v.3. for targeting upregulated upstream regulators.**

**Table S12. List of predicted antagonist drugs from DGIdb v.3. for targeting downregulated upstream regulators.**

## Notes

https://www.ncbi.nlm.nih.gov/bioproject/?term=PRJNA972323

### Competing Interest Statement

The authors have declared no competing interest.

### Summary of Updates

Figure 10 and associated text revised. The changes do not alter the overall conclusions of the research.

