## Supplementary figures and images for "A transcriptomics-based drug repositioning approach to identify drugs with similar activities for the treatment of muscle pathologies in spinal muscular atrophy (SMA) models"

### Supplementary Figure 1

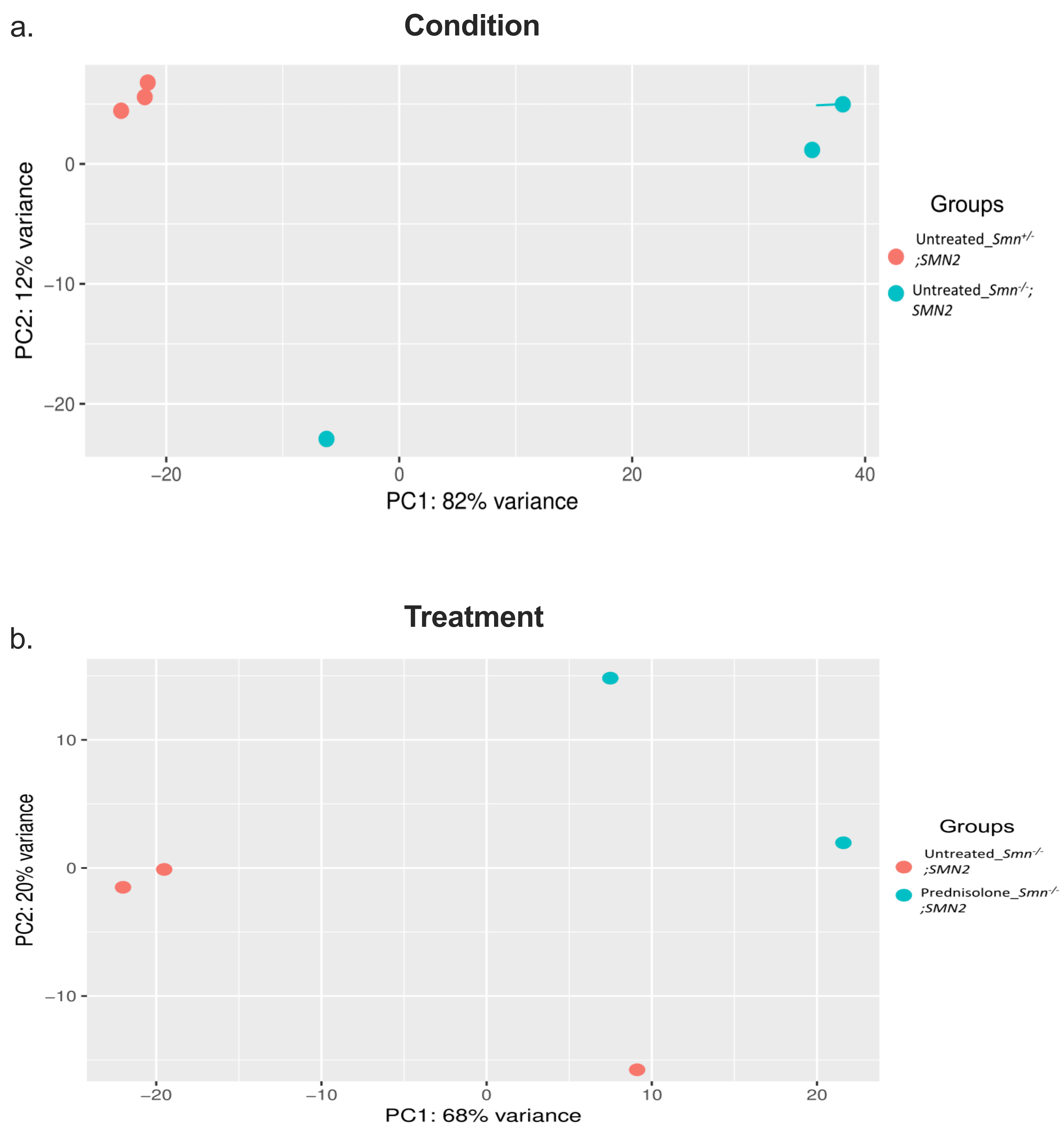

### Supplementary Figure 2

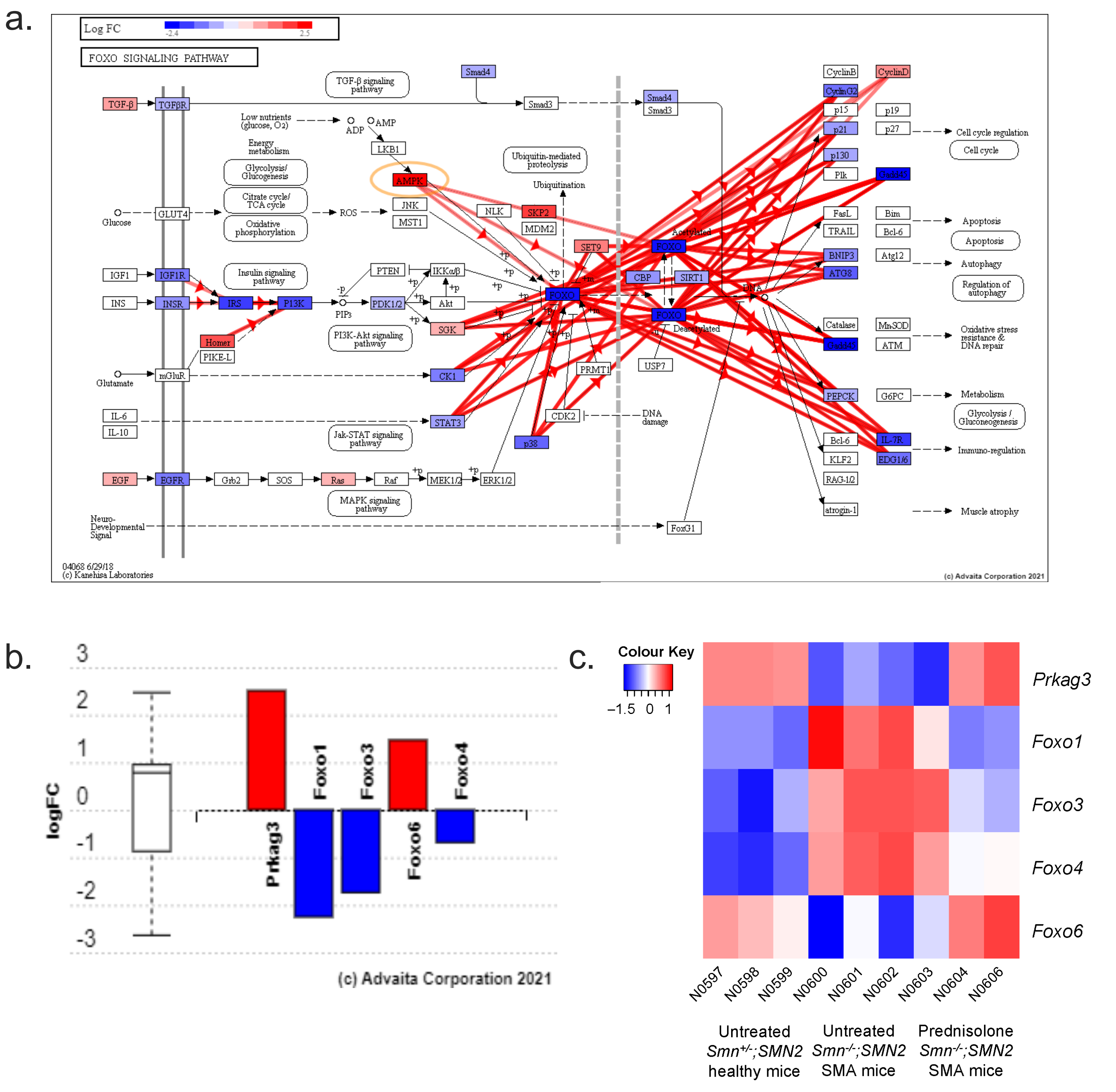

### Supplementary Figure 3

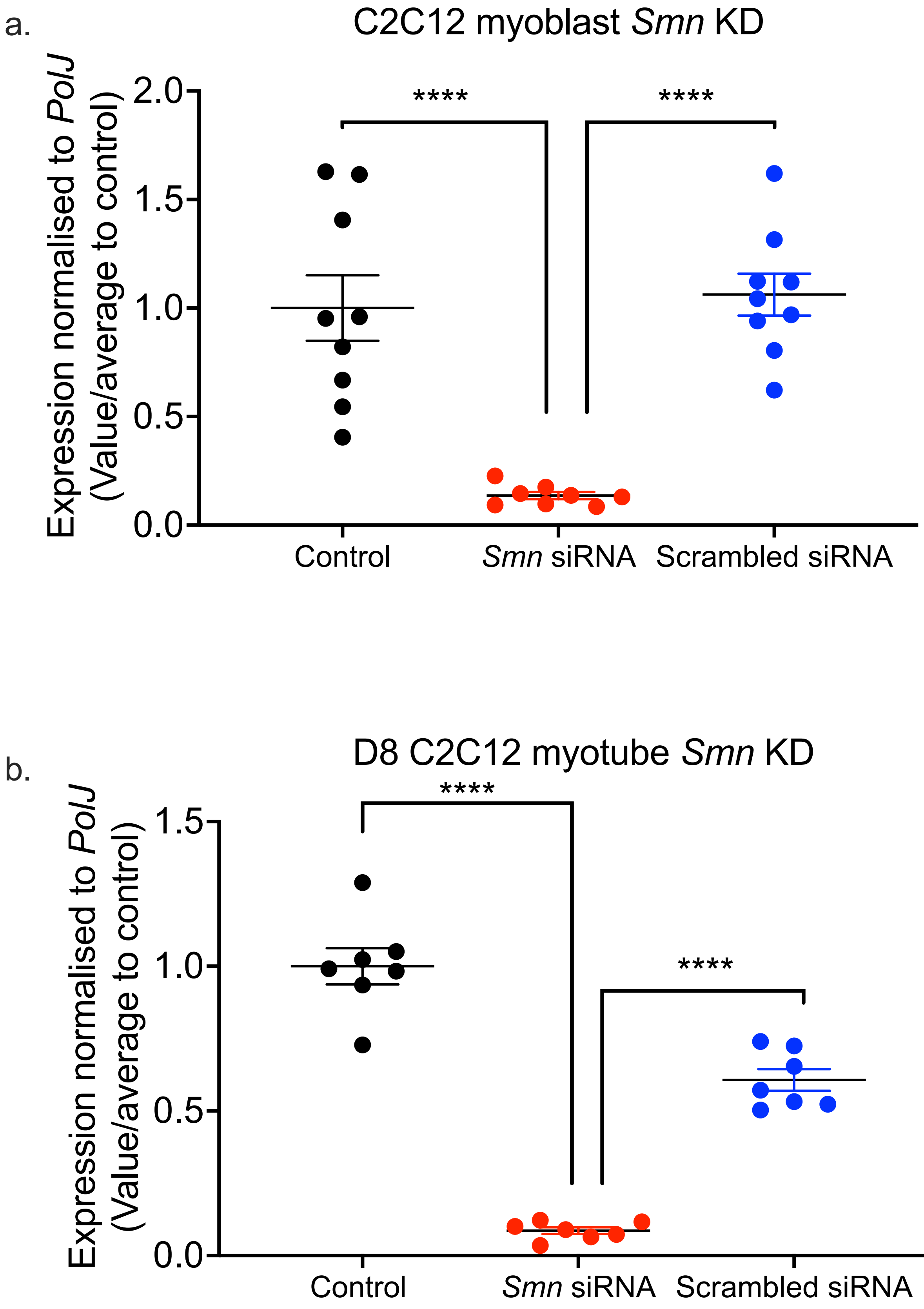

### Supplementary Figure 4

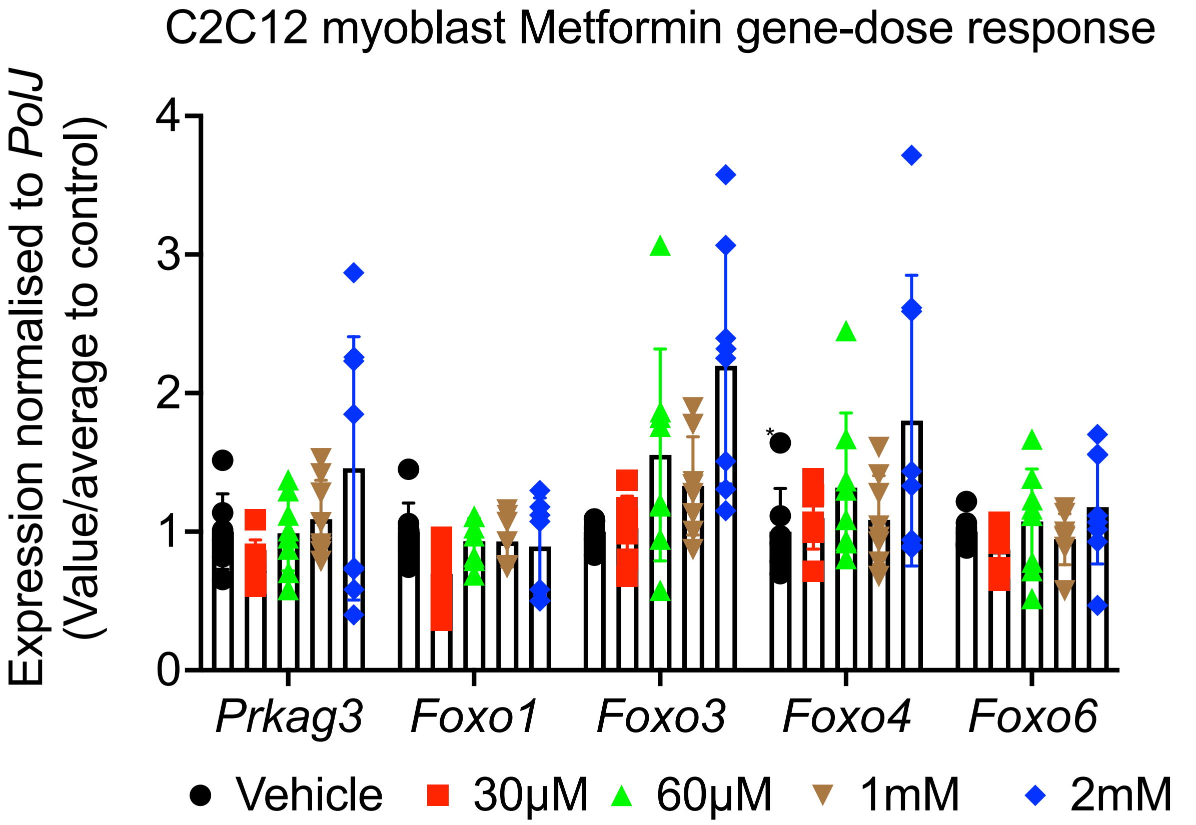

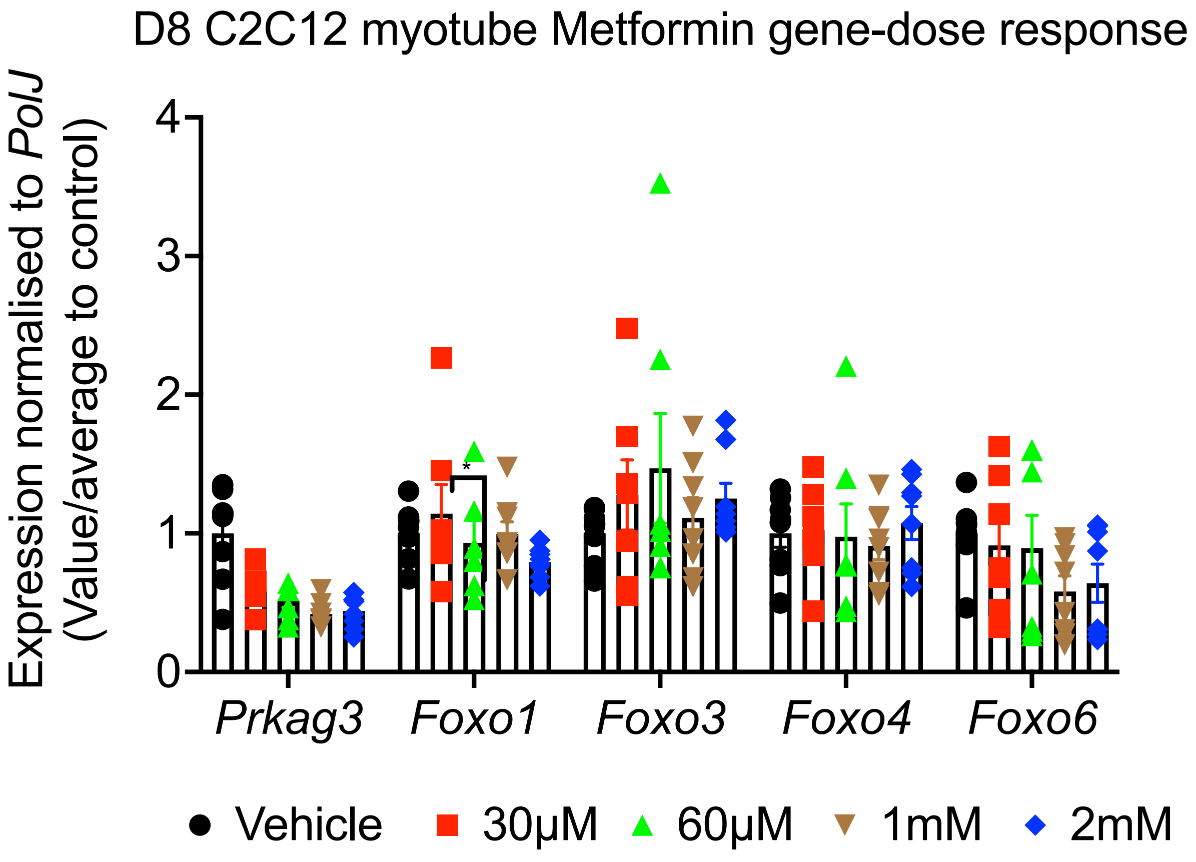

### Supplementary Figure 5

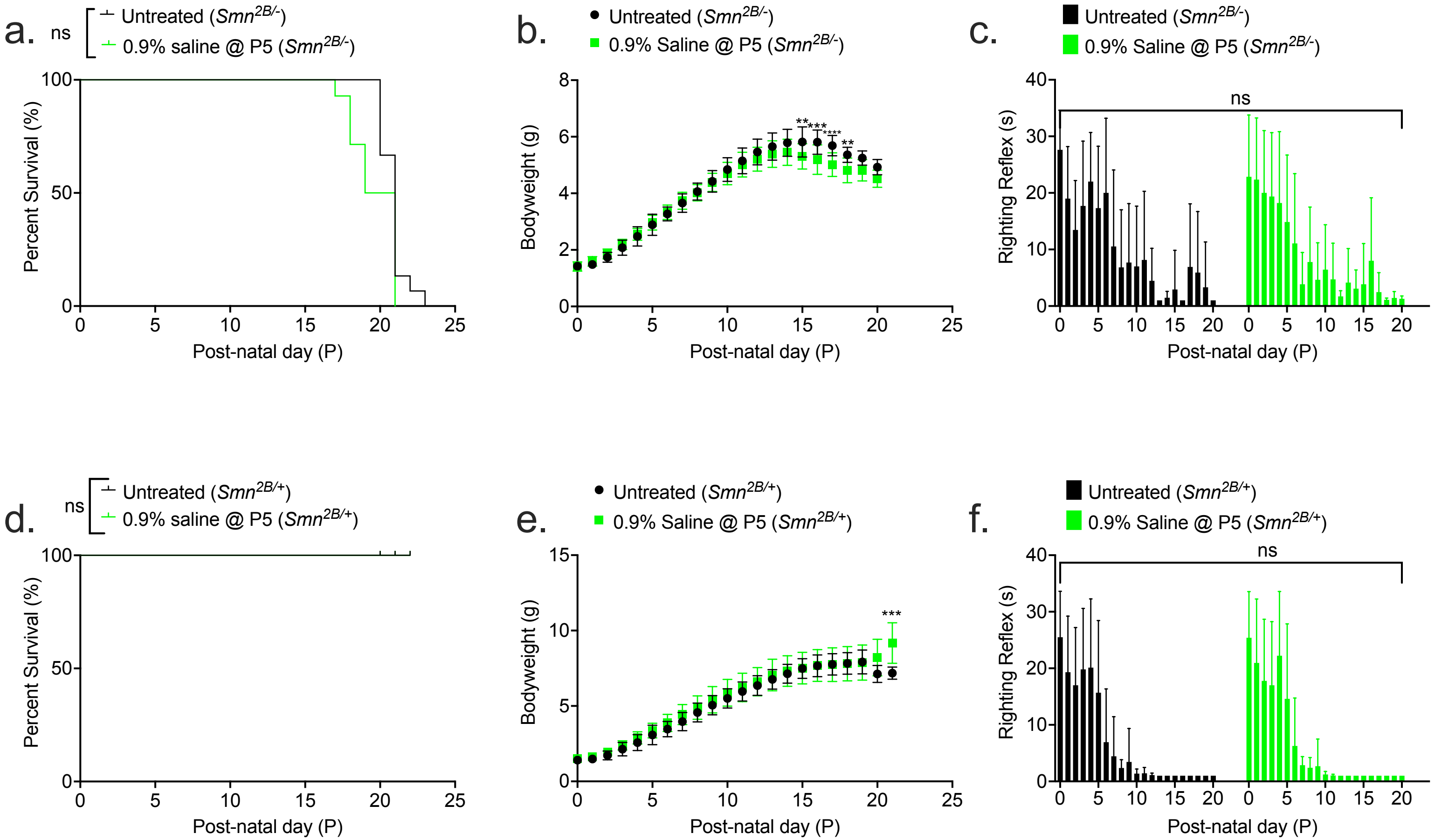

### Supplementary Figure 6

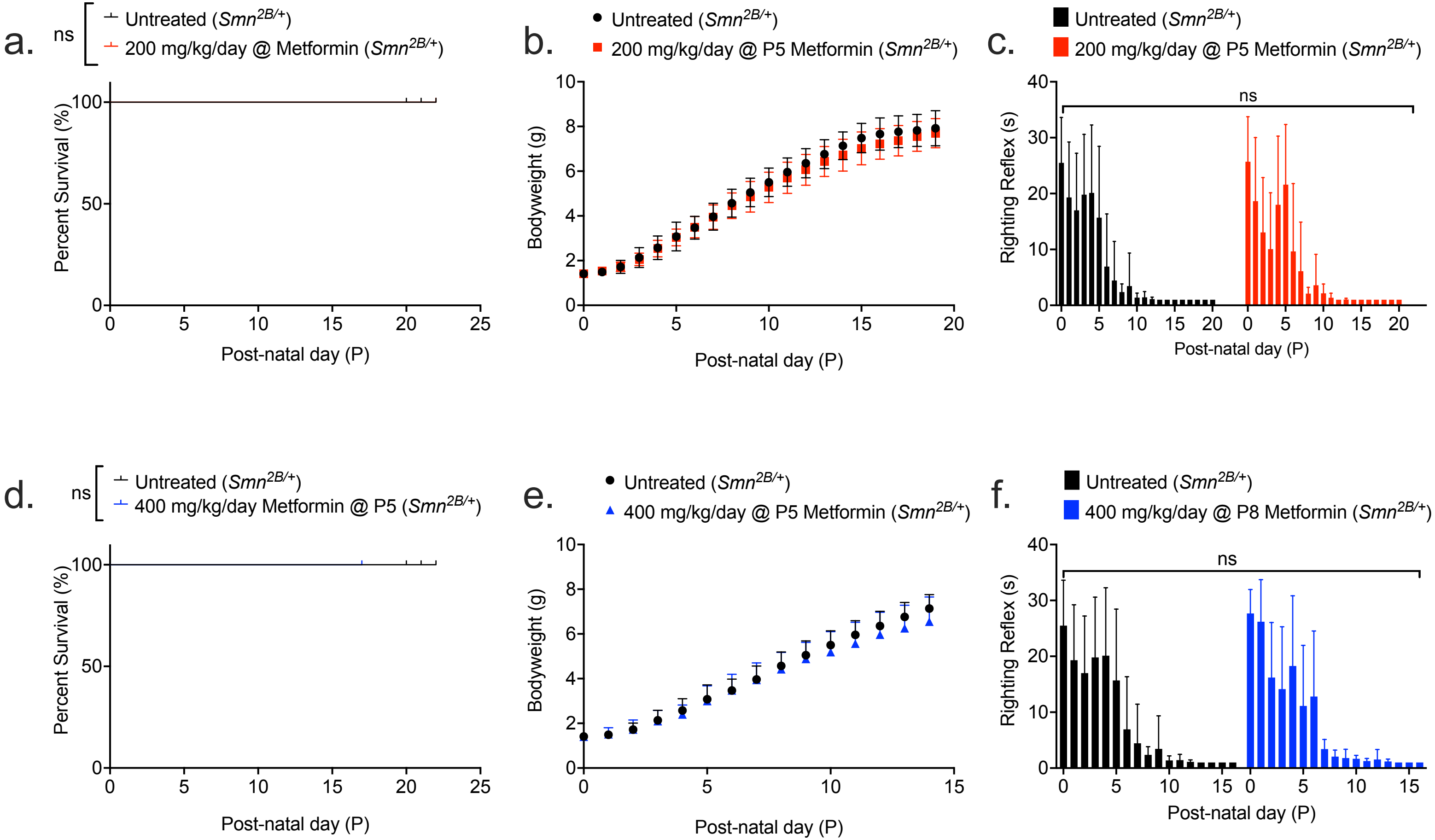

### Supplementary Figure 7

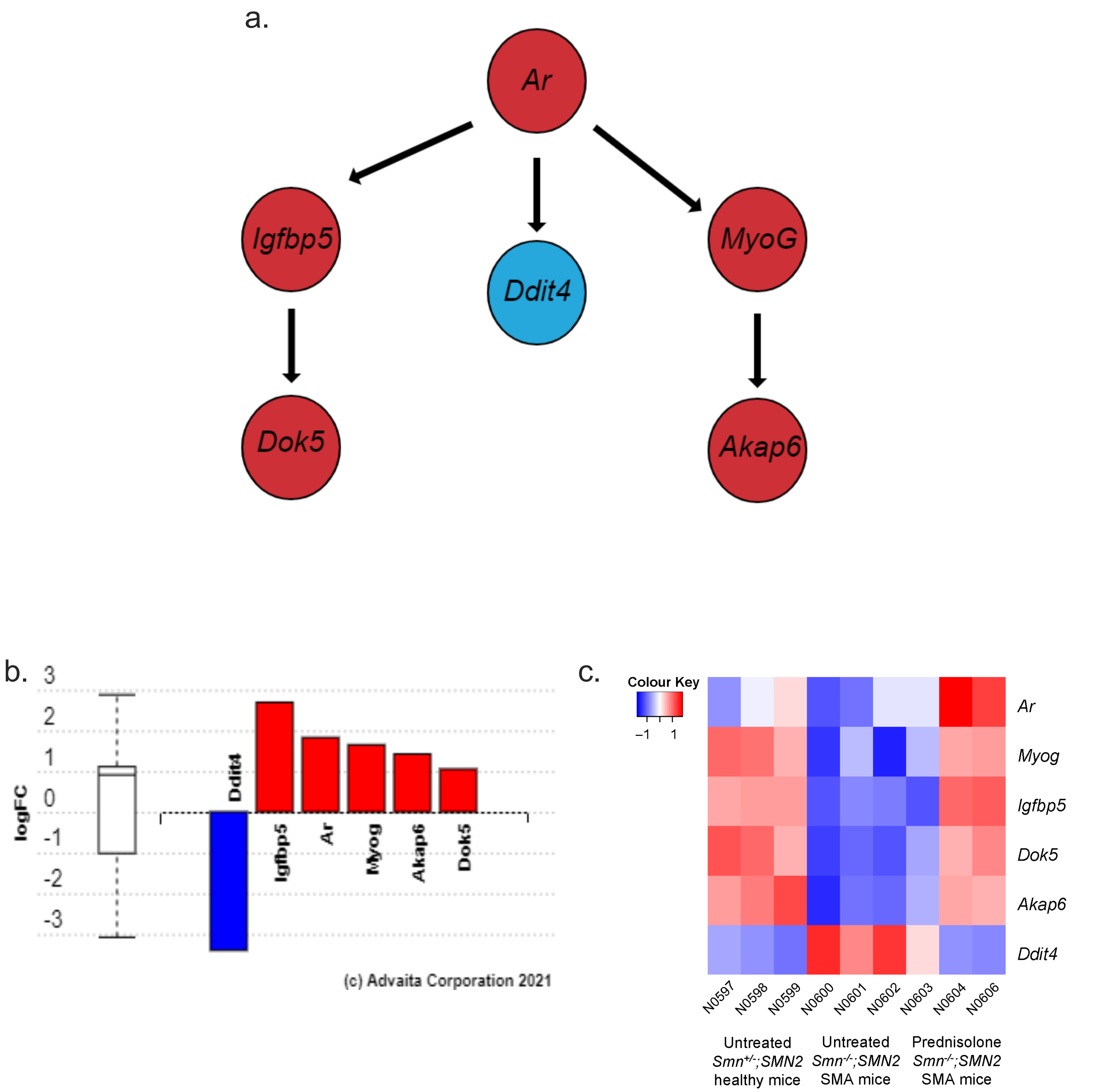

### Supplementary Figure 8

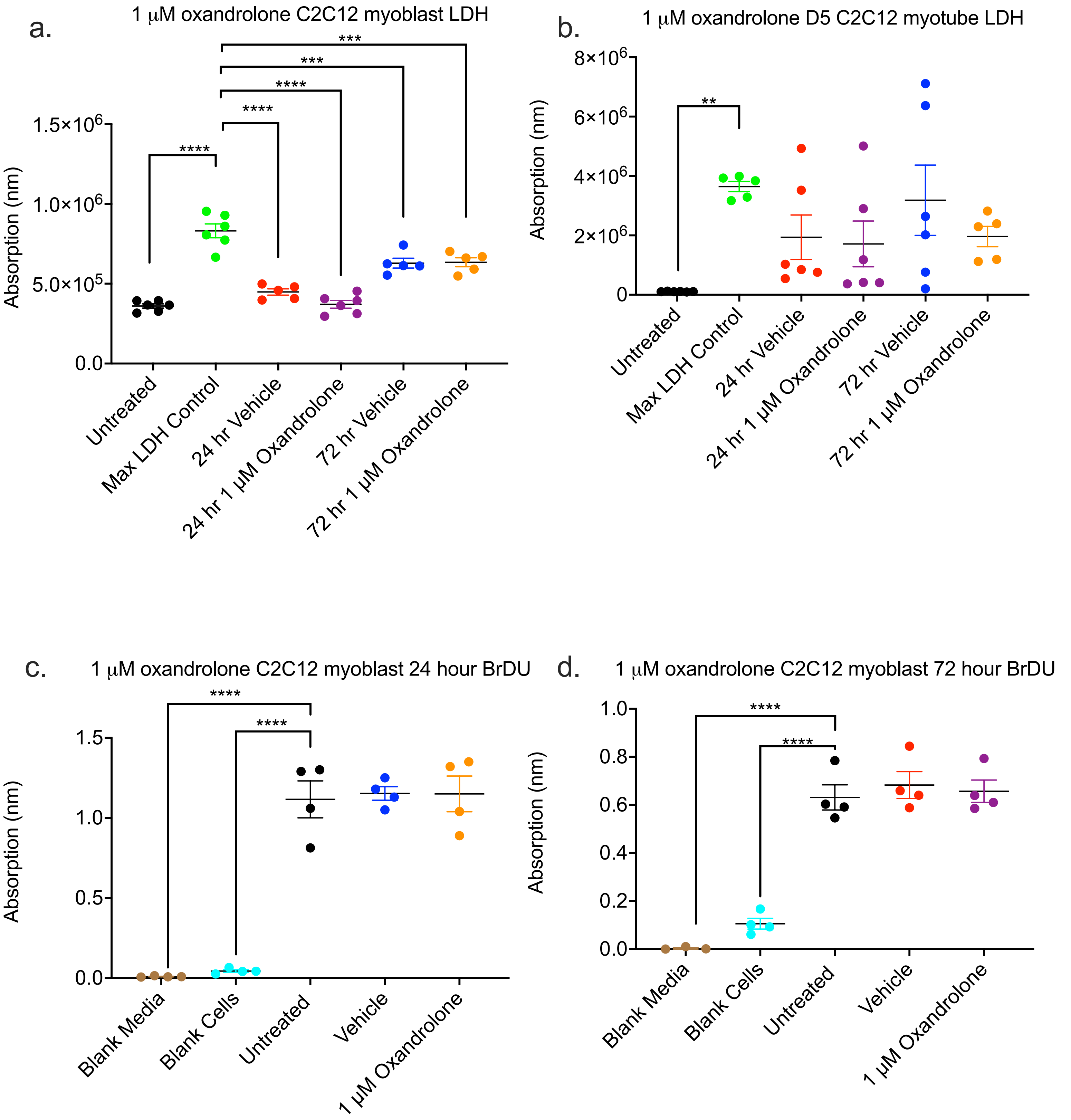

### Supplementary Figure 9

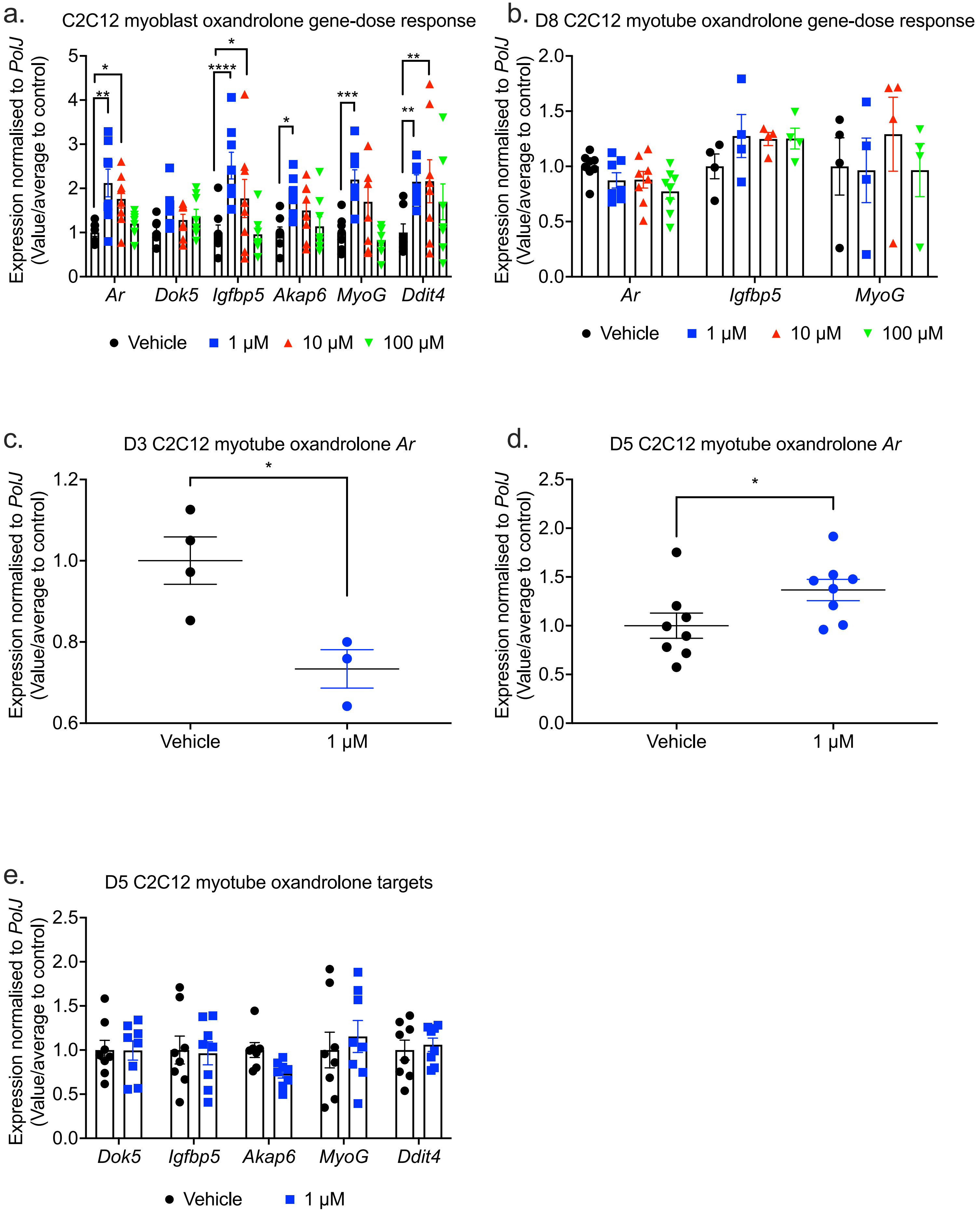

### Supplementary Figure 10

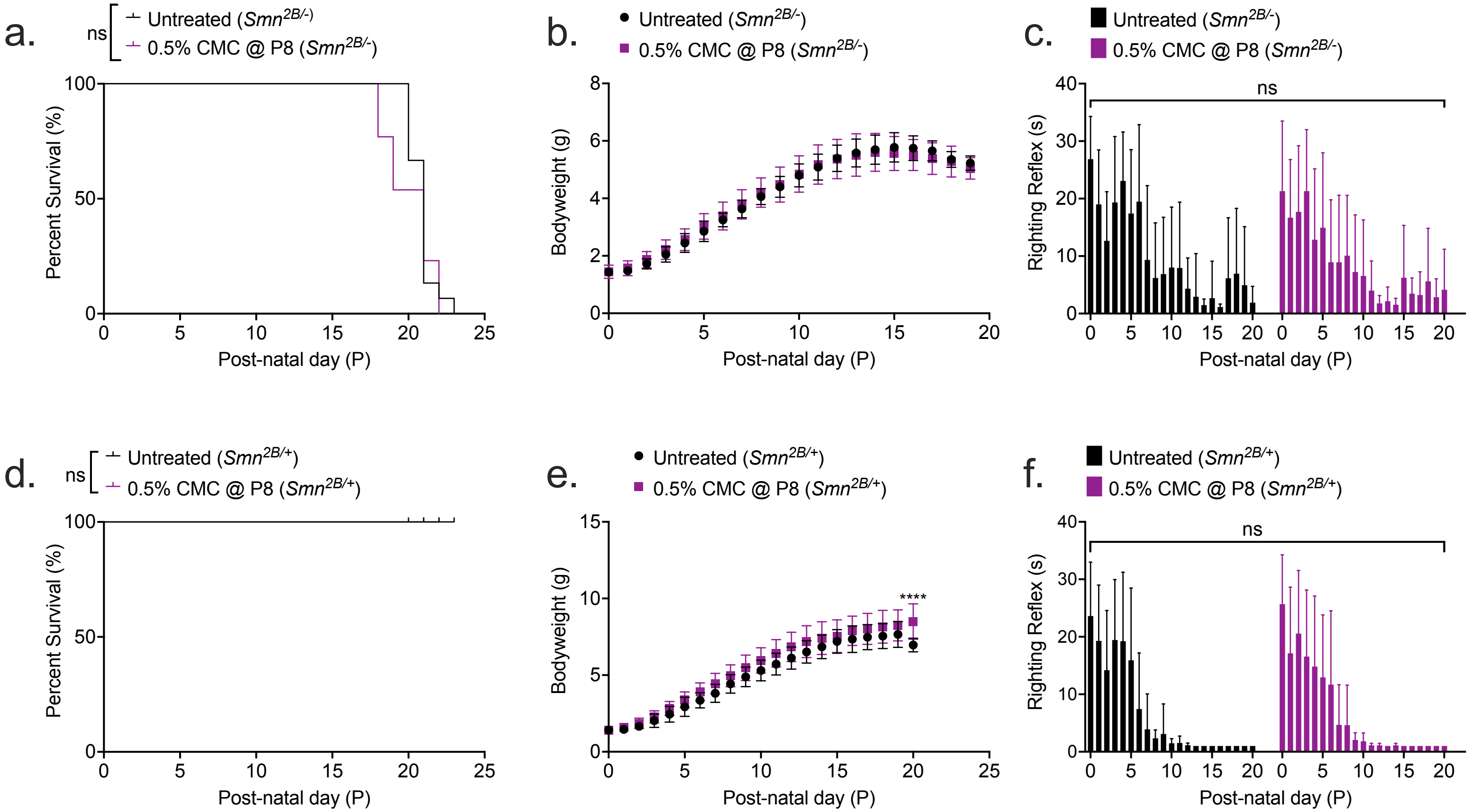

### Supplementary Figure 11

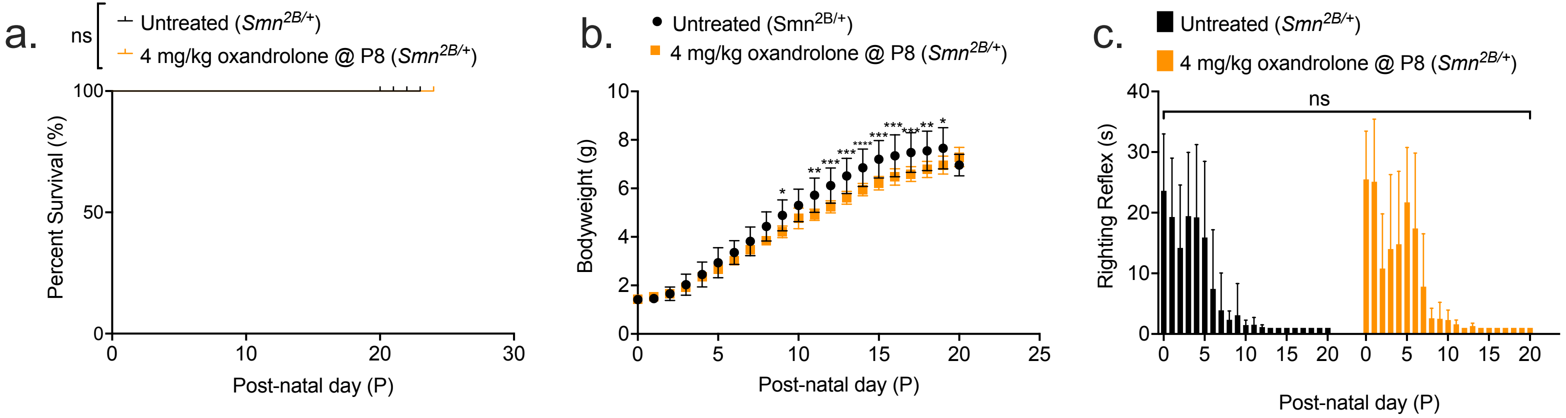
