## Supplementary Table 1 for "A transcriptomics-based drug repositioning approach to identify drugs with similar activities for the treatment of muscle pathologies in spinal muscular atrophy (SMA) models"

| **Table S1. RNA sequencing sample groups.** | | | |
| --- | --- | --- | --- |
| **Untreated**  ***Smn^+/-^;SMN2* mice** | **Untreated**  ***Smn^-/-^;SMN2* mice** | **Prednisolone-treated**  ***Smn^-/-^;SMN2* mice** | **Prednisolone-treated *Smn^+/-^;SMN2* mice** |
| N0597  N0598  N0599 | N0600  N0601  N0602 | N0603  N0604  N0606 | N0605  N0607 |
