## Supplementary Table 2 for "A transcriptomics-based drug repositioning approach to identify drugs with similar activities for the treatment of muscle pathologies in spinal muscular atrophy (SMA) models"

| **Table S2. Murine primers for qPCR.** | | |
| --- | --- | --- |
| **Genes** | **Forward Primers (5’ – 3’)** | **Reverse Primers (5’ – 3’)** |
| *PolJ* | ACC ACA CTC TGG GGA ACA TC | CTC GCT GAT GAG GTC TGT GA |
| *Smn* | TGC TCC GTG GAC CTC ATT TCT T | TGG CTT TCC TGG TCC TAA TCC TGA |
| *Atrogin-1* | TCA AAG GCC TCA CGA TCA CC | CCT CAA TGA CGT ATC CCC CG |
| *MuRF-1* | GAG AAC CTG GAG AAG CAG CT | CCG CGG TTG GTC CAG TAG |
| *Prkag3* | GGT CAT CTT TGA CAC GTT | AGA GGA GCT GCC CTC ACA C |
| *FoxO1* | CTA CGA GTG GAT GGT GAA GAG C | CCA GTT CCT TCA TTC TGC ACT CG |
| *FoxO3* | GGA AGG GAG GAG GAG GAA TG | CTC GGC TCC TTC CCT TCA G |
| *FoxO4* | CAA GAA GAA GCC GTC TGT CC | CTG ACG GTG CTA GCA TTT GA |
| *FoxO6* | AGA GCG CCC CGG ACA AGA GA | GCC GAA TGG AGT TCT TCC AGC C |
| *Ar* | TAC CAG CTC ACC AAG CTC CT | GAT GGG CTT GAC TTT CCC AG |
| *MyoG* | GTG TAA GAG GAA GTC TGT GTC GG | GCT CAA TGT ACT GGA TGG CG |
| *Igfbp5* | AAG AGC TAC GGC GAG CAA ACC A | GCT CGG AAA TGC GAG TGT GCT T |
| *Akap6* | AAG GAA CGA GCG CCG AGA AAC A | TGC TGG CAC AAC CTC AGA ATG G |
| *Dok5* | CGT TAT GGA CGA GAC ACC ACG T | GAG TGG ACC TTC TGG TAG ATG G |
| *Ddit4* | CCT GCG CGT TTG CTC ATG CC | GGC CGC ACG GCT CAC TGT AT |
| *Hk2* | GAA GGG GCT AGG AGC TAC CA | CTC GGA GCA CAC GGA AGT T |
| *Glut4* | GAC GGA CAC TCC ATC TGT TG | CAT AGC TCA TGG CTG GAA CC |
| *Pgc1-α* | TGG AGT GAC ATA GAG TGT GCT GC | CTC AAA TAT GTT CGC AGG CTC A |
| *Nrf1* | CAG CAC CTT TGG AGA ATG TG | CCT GGG TCA TTT TGT CCA CA |
| *Tfam* | CAA GTC AGC TGA TGG GTA TGG | TTT CCC TGA GCC GAA TCA TCC |
| *Nfdus1* | GTG GAT GCT GAA GCC TTA GTA GC | GGA ACG TAA GTC TGT ACC AGC TC |
